# CD4^+^ T cells display a spectrum of recall dynamics during re-infection with malaria parasites

**DOI:** 10.1101/2023.03.02.530907

**Authors:** Hyun Jae Lee, Marcela L. Moreira, Shihan Li, Cameron G. Williams, Oliver P. Skinner, Saba Asad, Takahiro Asatsuma, Michael Bramhall, Zhe Jiang, Jessica A. Engel, Megan S. F. Soon, Jasmin Straube, Irving Barrera, Evan Murray, Fei Chen, Jason Nideffer, Prasanna Jagannathan, Ashraful Haque

## Abstract

Children in malaria-endemic regions can experience multiple *Plasmodium* infections over a short period of time, with *in vitro* CD4^+^ T cell recall responses becoming more regulatory with increasing age and exposure. This suggests that repeated infection qualitatively changes CD4^+^ T cells, although the heterogeneity and dynamics of these responses await systematic analysis *in vivo*. Here, we examined TCR transgenic PbTII and polyclonal CD4^+^ T cells during *Plasmodium* re-infection in mice, in conjunction with scRNA-seq/TCR-seq and spatial transcriptomics at near single-cell resolution. PbTII cells gave rise to multiple antigen-experienced states in different areas of the spleen after primary infection and antimalarial treatment, including ongoing GC responses and T-cell zone memory. Upon re-infection, Th1-memory PbTII cells initiated a rapid effector response prior to proliferating, while GC Tfh cells of the same antigen specificity were entirely refractory within the same organ. Transcriptome dynamic modelling and network analysis of Th1 recall revealed a biphasic wave of RNA processing that firstly preceded immune effector transcription, and later accompanied cellular proliferation. Importantly, Th1 recall constituted a partial facsimile of primary Th1 responses, with no unique genes amongst the small subset of those upregulated upon re-infection. Finally, we noted a similar spectrum of antigen-experienced states and recall dynamics by polyclonal CD4^+^ T cells with diverse TCRs. Therefore, during re-infection with *Plasmodium*, persisting GC Tfh cells remained unaltered transcriptionally, Tcm/Tfh-like cells exhibited minimal proliferation, and Th1-memory cells displayed a rapid, proliferating IL-10-producing Tr1 response consistent with a shift towards immune-regulation. These data highlight a broad spectrum of simultaneous CD4^+^ T cell responses that occur in the spleen during re-infection with malaria parasites.

**Highlights:**

- Splenic TCR transgenic CD4^+^ T cells are highly heterogeneous prior to re-infection.
- Persisting GC Tfh cells are refractory to re-activation during re-infection.
- Th1-memory cells rapidly upregulate RNA processing prior to effector function and proliferation.
- Th1-recall is an imperfect but faithful facsimile of primary Th1 responses.
- A spectrum of recall states is observed in polyclonal CD4^+^ T cells with diverse TCRs.

## Introduction

Malaria is a mosquito-borne disease caused by infection *Plasmodium* parasites, with 247 million cases and 619,000 deaths in 2021^1^. It remains a major global health burden, with malaria morbidity and mortality highest amongst young children^1^. In malaria endemic regions, it is common for children to experience repeated infections over a short timeframe of weeks to months^2, 3^. Cumulative infections eventually result in acquisition of clinical immunity against malaria infection where children are protected from severe malaria syndromes, are less likely to develop febrile symptoms, and can tolerate low to intermediate levels of parasitemia^4, 5^.

CD4^+^ T cells orchestrate coordinated immune responses to pathogens and likely play an important role in facilitating immunity to malaria. Assessment of parasite-specific CD4^+^ T cells in circulating blood, via antigen-specific *in vitro* restimulation and intracellular-cytokine staining, revealed an increased propensity for interleukin-10 production with increased age and exposure^6, 7^. This suggested that repeated infection may alter CD4^+^ T cells qualitatively. Given that parasite-specific, isotype-switched antibodies accumulate in the bloodstream^8, 9^, it is likely that germinal centre T follicular helper (GC Tfh) cell responses occur in these children, although direct visualisation in the spleen is currently challenging. How multiple types of antigen-experienced CD4^+^ T cell respond during repeat infection with malaria parasites remains unknown. More generally, our understanding of CD4^+^ T cell memory recall is focussed on effector memory cells, and their expression of a handful of molecules including cytokines IFN-γ, TNF, IL2, IL10, costimulatory molecules such as OX40, and proliferative markers, *e.g.* Ki67 and BrdU incorporation. The broader relationship between primary Th1 and recall Th1 responses is unknown. Moreover, the dynamics of transcriptional change exhibited during recall remains unclear.

Previously, we employed TCR-transgenic (PbTII) CD4^+^ T cells, specific for an epitope from *Plasmodium* Heat Shock Protein 90^10, 11^, to map the transcriptional dynamics that underlie clonal expansion, Th1/Tfh effector fate choice^12, 13^, and transit to memory or exhausted states in experimental primary malaria^14^. These studies suggested roles for inflammatory monocytes and B cells, and the genes *Tcf7* and *Lgals1* in controlling T helper 1 (Th1)/T follicular helper (Tfh) fate choice, as well as revealing that memory emerges gradually over a 3-4 week period from effector counterparts. Importantly, droplet-based scRNA-seq at late timepoints suggested a complex landscape of cellular states amongst splenic PbTII cells comprised of Th1 effector memory cells, T central memory (Tcm) cells, germinal centre (GC) Tfh cells, Tfh cells, and a range of other mixed phenotypes. Whether and how these transcriptionally diverse cells responded during a second infection remained unclear. Finally, the relevance of TCR transgenic PbTII cells to polyclonal CD4^+^ T cell recall remained untested.

Here, we map over time splenic *in vivo* responses of antigen-experienced, TCR-transgenic and polyclonal CD4^+^ T cells during a second malaria infection, using a combination of droplet-based scRNA-seq, VDJ-seq, spatial transcriptomics at near-single-cell resolution, and computational modelling. We define CD4^+^ T cell heterogeneity before and after re-infection, reveal differential recall dynamics, and predict novel biological processes and gene markers that accompany recall during a second exposure to blood-stage malaria parasites.

## Results

### *Plasmodium-*specific CD4^+^ T cells exhibit a spectrum of transcriptional states in the spleen after primary infection and drug treatment

To map CD4^+^ T cell recall responses during re-infection with malaria parasites, we first tested whether transfer of PbTII cells, followed by primary *Plasmodium chabaudi chabaudi* AS (*Pc*AS) infection and drug cure, consistently seeded the spleen with heterogeneous antigen-experienced *Plasmodium-*specific CD4^+^ T cells, as suggested by our previous study^14^. We recovered splenic PbTIIs from mice 4 weeks after primary *Pc*AS infection and antimalarial drug treatment (Extended Data Fig. 1A), and processed these via droplet-based scRNA-seq (Figure 1A). We removed poor-quality transcriptomes and computationally integrated with our previous dataset using *Seurat*^15^ to remove technical batch effects (Extended Data Fig. 1B), as indicated by Principal Component Analysis (principal components 1-6) and Uniform Manifold Approximation and Projection (UMAP) (Figure 1B; Extended Data Fig. 1C). This revealed similar proportions of cells from our two independent experiments in each transcriptionally-defined cluster (Extended Data Fig. 1D), also noted using an alternative integration approach, *Harmony*^16^ (Extended Data Fig. 1E). This revealed that PbTII cells adopted a consistent heterogeneous set of transcriptional states after primary infection and drug cure. Assessment of canonical naïve/Tcm (*Ccr7*, *Sell*, *Tcf7*, and *Klf2*), helper (*Cxcr6*, *Ifng*, *Cxcr5,* and *Bcl6*), proliferative (*Mki67*), and Type I IFN-associated (*Ifit1*, *Ifit3*, and *Irf7*) genes (Figure 1C), supported the existence of GC Tfh (marked by high expression of *Cxcr5* and *Bcl6*), Tfh, Th1-effector memory, Tcm, IFN-associated (IFN-high) and proliferating states within PbTII cells (Figure 1D). However, many of these states appeared not as distinct clusters, but as a transcriptional continuum (Figure 1D). Thus, after primary infection and drug cure, a complex landscape of transcriptomic states was exhibited by splenic PbTII cells, primarily consisting of memory-like and Tfh-like states.

**Figure 1:**
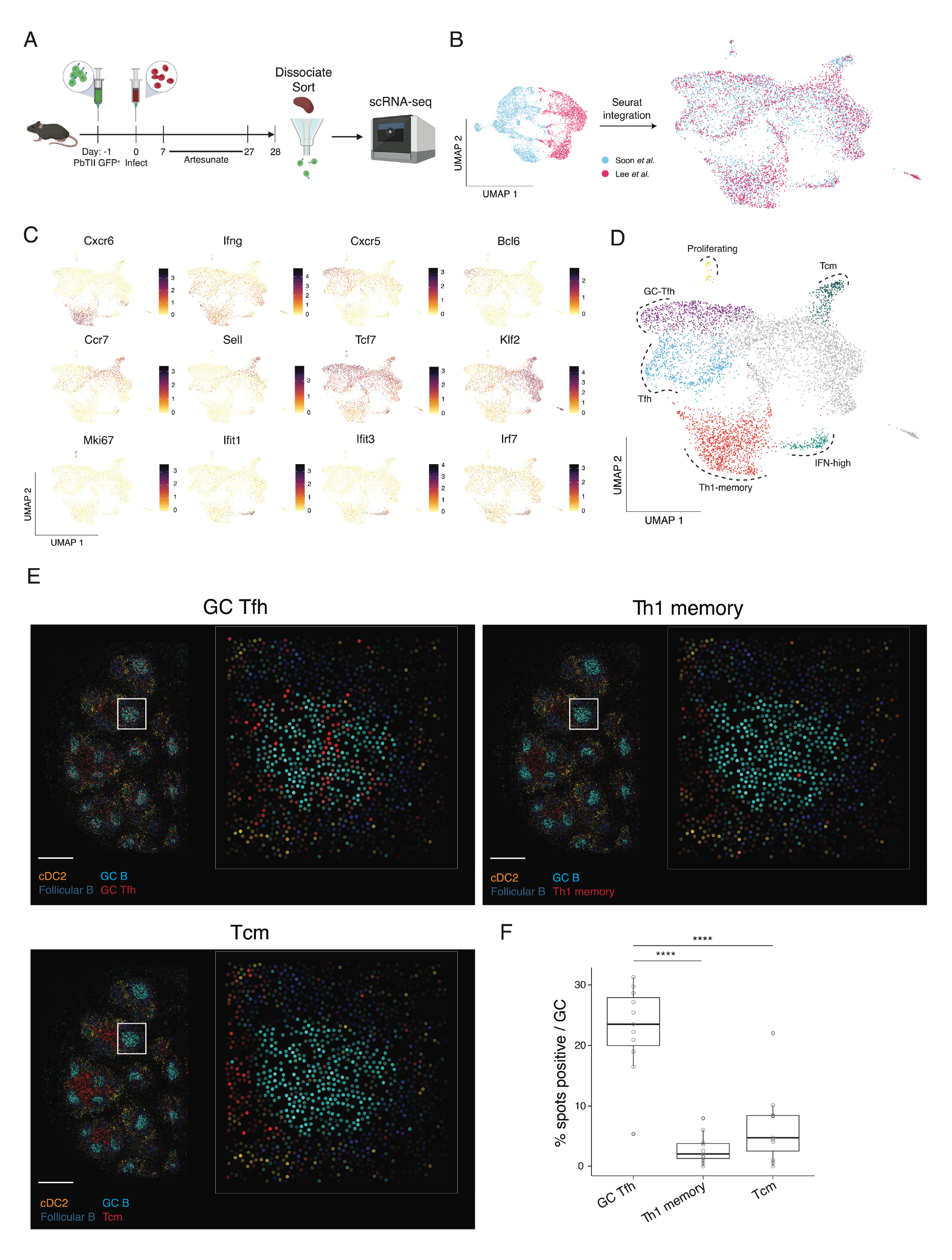
PbTII cells exhibit a spectrum of transcriptional states in the spleen after primary infection and antimalarial drug treatment. **(A)** Schematic of scRNA-seq experiment to study PbTII cells at day 28 post-infection. **(B)** UMAPs of PbTII cell-derived scRNA-seq data before and after integration of our two independent experiments, this study and Soon et al.^14^ - cells coloured by experimental origin. **(C)** UMAPs of PbTII cell gene expression patterns for Th1 (*Cxcr6*, *Ifng*), Tfh (*Cxcr5*, *Bcl6*), Tcm (*Ccr7*, *Sell*, *Tcf7*, *Klf2*), proliferation (*Mki67*), and IFN-high (*Ifit1*, *Ifit3*, *Irf7*) genes. **(D)** UMAP of PbTII cell states depicted by colours and dotted boundaries, with those belonging to no distinct cell state coloured grey. **(E)** Estimated cell-type abundance for GC Tfh, Tcm, and Th1 memory (red), compared with the same cDC2, GC B cells and follicular B cells in each panel, assessed across a *Slide-seqV2* spatial transcriptomic map of mouse spleen at day 30 p.i. - scale bar: 500µm: white boxes shown at higher magnification within each inset panel, with each dot being a 10µm bead **(F)** Box plots showing % positive occurrence of GC Tfh, Tcm, or Th1-memory cells in each GC. Cell abundance (computed using *cell2location*) greater than 0.2 is considered a positive occurrence. For box plot, centre line indicates median and box limits upper and lower quartiles. Statistical analysis performed using Welch two sample t-test between each cell type. \*\*\*\**P*<0.0001.

To determine if cells transcriptionally annotated as GC Tfh were indeed located in GC B-cell regions of the spleen, we conducted spatial transcriptomic assessment of spleen sections using *Slide-seqV2*, a technique offering near single-cell resolution^17, 18^. As expected, discrete GC B cell transcriptomic regions were noted within B cell follicles in infected mice (Figure 1E; Extended Data Fig. 2A-C). Importantly, GC Tfh transcriptomic signals derived from PbTII data, were enriched in GC regions compared to either Th1 or Tcm transcriptomes, which instead were confined to areas between B cell follicles, inferred to be T-cell zones enriched with cDC2 cells (Figure 1F). These data suggested *Plasmodium-*specific CD4^+^ T cells exhibited GC Tfh states that co-localised with GC B cells 28 days after infection and drug cure, as well as circulating memory states residing in T-cell zones. Thus, a variety of CD4^+^ T cell memory and Tfh states co-existed in distinct areas of the spleen after primary *Plasmodium* infection and anti-malarial drug cure.

**Figure 2:**
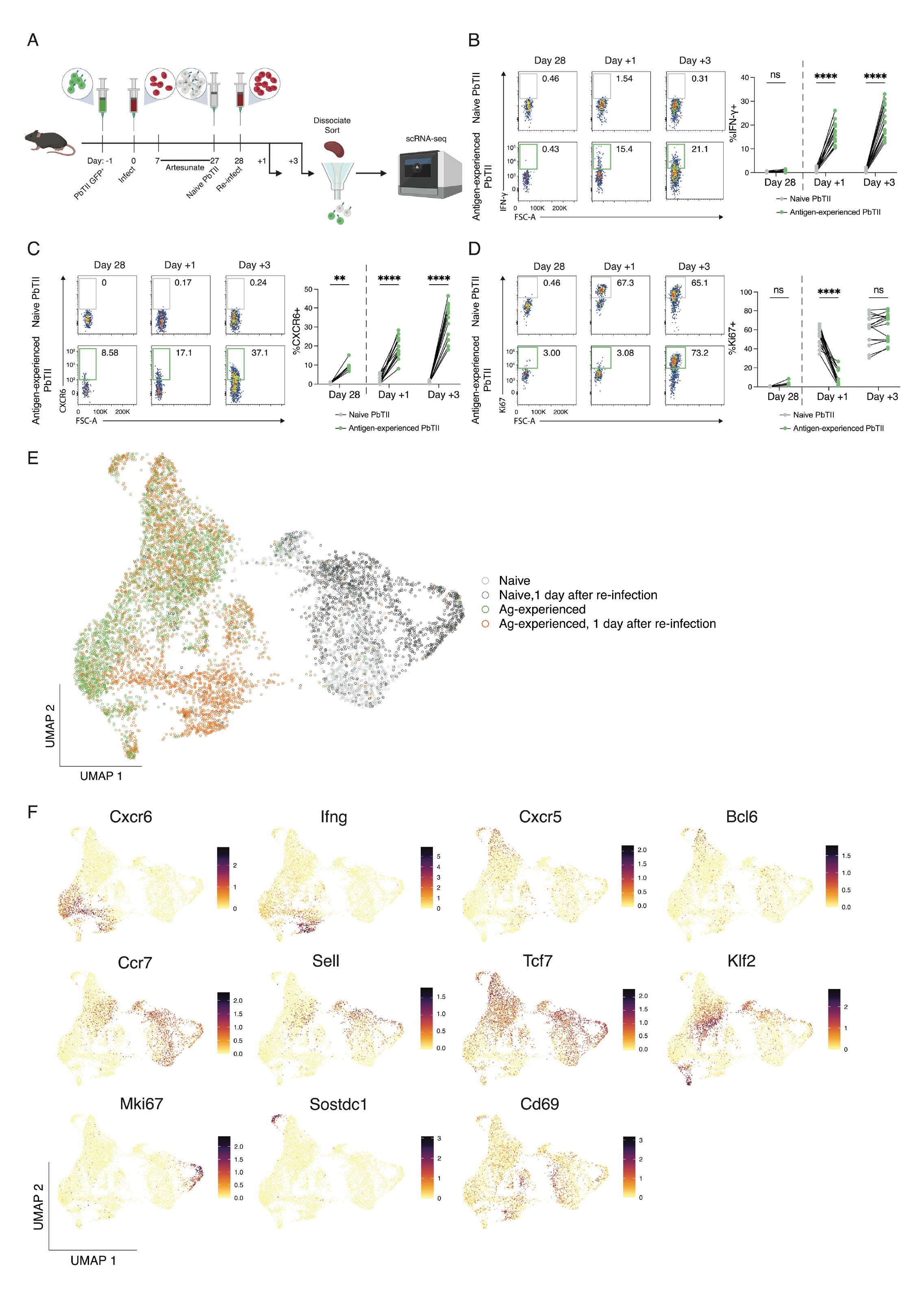
Re-infection triggers an early transcriptional response from Th1-memory PbTIIs. **(A)** Schematic of scRNA-seq experiment to study *in vivo* responses of antigen-experienced versus naïve PbTII cells during re-infection. **(B/C/D)** Representative FACS plots showing **(B)** IFNγ, **(C)** CXCR6, and **(D)** Ki67 expression in antigen-experienced PbTII cells at day 28 p.i. and 1 and 3 days post re-infection, compared to co-transferred naïve comparator PbTIIs. Data combined from two independent experiments showing similar results (n=9-15 mice) - statistical analysis performed using 2-way ANOVA. *p<0.05, ****p<0.0001. **(E)** UMAP of antigen-experienced PbTII cells and naïve comparators, prior to and 1 day after re-infection. **(F)** UMAPs of PbTII expression of various genes associated with Th1 (*Cxcr6*, *Ifng*), Tfh (*Cxcr5*, *Bcl6*), Tcm (*Ccr7*, *Sell*, *Tcf7*, *Klf2*), proliferation (*Mki67*), *Sostdc1*^+^ cells, and early activation (*Cd69*).

### Antigen-experienced CD4^+^ T cells display a variety of recall dynamics during re-infection

To map responses of heterogeneous splenic CD4^+^ T cells during re-infection, we first determined by flow cytometry whether antigen-experienced PbTIIs could mount a recall response. Our previous study indicated a minority (∼20-30%) of PbTIIs secreted IFN-γ without proliferating within 24 hours of re-infection with homologous parasites^14^. Here, we extended these findings, confirming that in contrast to CTV-labelled naïve PbTIIs (Extended Data Fig. 3) co-transferred one day prior to re-infection (Figure 2A), 10-30% of antigen-experienced PbTIIs (Extended Data Fig. 3) rapidly secreted IFN-γ and upregulated CXCR6 by day 1 to day 3 of re-infection (Figure 2B and 2C), consistent with a pronounced Th1-recall response. In addition, the majority of PbTIIs, but not all, had upregulated Ki67 expression by day 3, suggesting a heterogeneous capacity for CD4^+^ T cell proliferation within three days of re-infection (Figure 2D). Together, these data indicated that a proliferative and a Th1 recall response were mounted by some but not all antigen-experienced PbTIIs.

**Figure 3:**
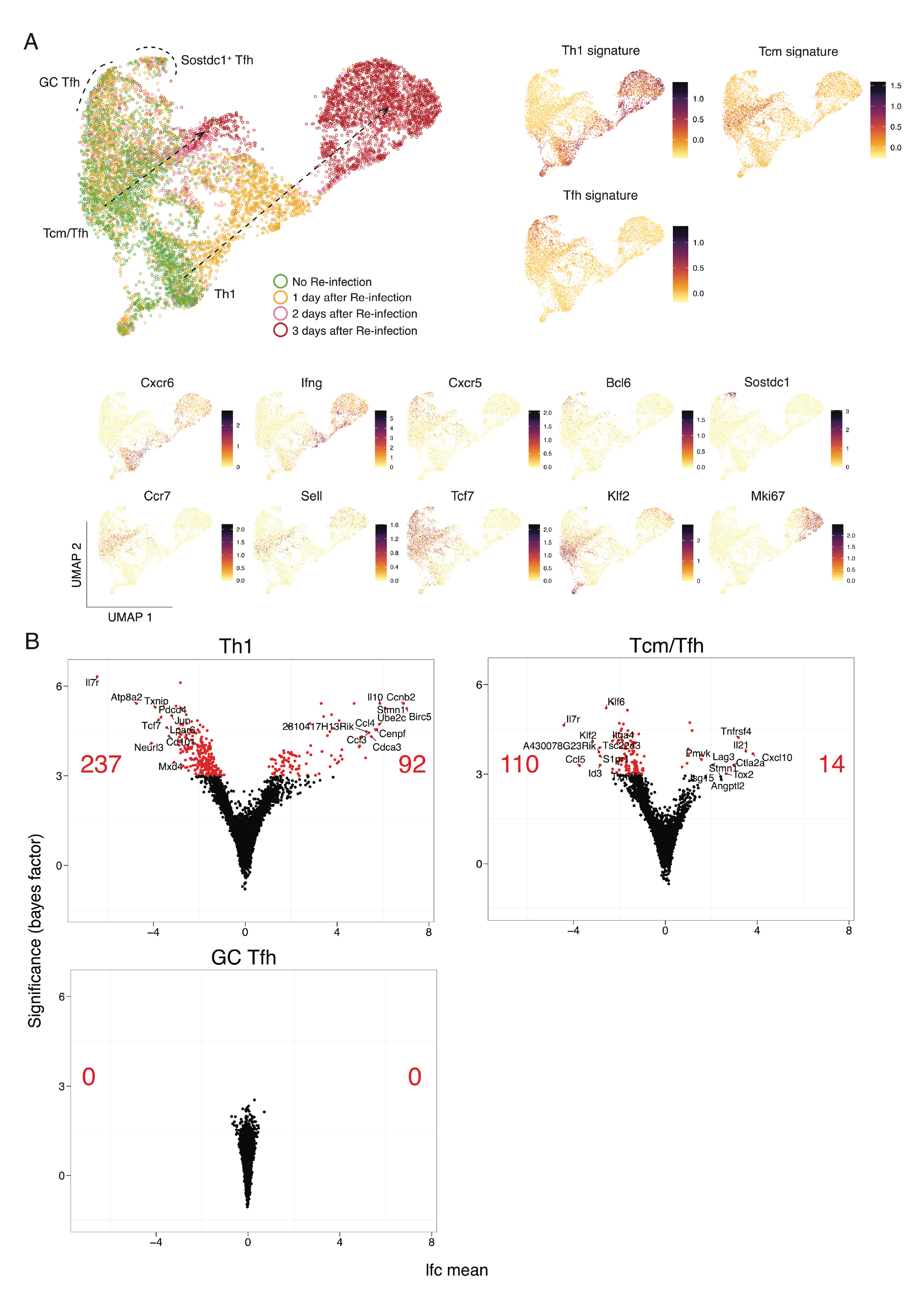
PbTII cells exhibit a spectrum of recall dynamics during re-infection. **(A)** (*Left*) UMAP representation of antigen-experienced PbTII cells prior to and 1, 2, and 3 days post re-infection; two apparent trajectories indicated by arrows; GC Tfh cells and *Sostdc1*^+^ Tfh cells marked with dotted boundaries. (*Right*) Th1, Tcm, and Tfh signature scores. (*Bottom*) Expression patterns for various genes associated with Th1 (*Cxcr6*, *Ifng*), Tfh (*Cxcr5*, *Bcl6*), *Sostdc1*^+^ cells, Tcm (*Ccr7*, *Sell*, *Tcf7*, *Klf2*), and proliferation (*Mki67*) states. **(B)** Volcano plots depicting the number and LogFC (lfc mean) of differentially expressed genes (genes with bayes factor > 3) from comparing Th1 cells (Left), Tcm/Tfh cells (Right), and GC Tfh cells (Bottom) prior to and post re-infection. Number of significantly upregulated/downregulated genes and the top 10 upregulated/downregulated genes annotated on volcano plots.

To test for heterogeneity in recall as suggested by flow cytometry, we compared antigen-experienced PbTIIs and naïve comparators by scRNA-seq 24 hours after re-infection (Figure 2E). Antigen-experienced cells enriched for Th1-markers *Cxcr6* and *Ifng* had indeed undergone transcriptomic change, suggestive of rapid Th1-recall (Figure 2F). In stark contrast, antigen-experienced PbTIIs enriched for *Tcf7* and either Tfh markers *Bcl6* and *Cxcr5*, or Tcm markers, *Sell, Ccr7* and *Klf2*, exhibited no transcriptomic change (Figure 2F). We also noted a minor population of GC Tfh-like cells that expressed high levels of *Sostdc1* (Figure 2F), previously implicated in promoting T follicular regulatory cells^19^. Finally, consistent with flow cytometric assessment, only naïve PbTII control cells upregulated *Mki67* over this timeframe (Figure 2F). Therefore, during the first 24 hours of re-infection, Th1-memory PbTIIs mounted a rapid non-proliferative effector recall response, while GC Tfh, Tfh, Tcm and *Sostdc1^+^* GC Tfh states remained transcriptionally unaltered. These data revealed striking heterogeneity in the early recall response of splenic, antigen-experienced CD4^+^ T cells of the same peptide-specificity.

To further test the apparent refractory nature of splenic Tfh and Tcm states relative to Th1-memory during re-infection, we extended our scRNA-seq analysis of antigen-experienced PbTIIs from day 1 to days 2 & 3 after re-infection, a time-period during which flow cytometry had suggested robust Th1 recall and emergence of a proliferative state. Low-dimensional UMAP embeddings after PCA, and annotation according to timepoint suggested three main trajectories, based on gene expression signatures and individual canonical genes, corresponding to Th1, Tcm and Tfh-like states (Figure 3A). Together, these suggested that Th1-recall dynamics were characterised by rapid immune effector expression followed by cellular proliferation by day 3. In contrast, GC Tfh and *Sostdc1^+^* GC Tfh remained unaltered over the three-day period. Tcm/Tfh-like cells displayed intermediate transcriptional change by day 3, notably devoid of *Mki67* upregulation, suggesting this cell state had not initiated proliferation (Figure 3A). Differential gene expression analysis for each inferred cell state prior to *versus* the peak of their apparent response revealed that antigen-experienced Th1-like cells differentially expressed 329 genes during re-infection, Tcm/Tfh-like cells 124 genes, and GC Tfh cells no genes (Figure 3B; Supplementary Tables 1-3). Together these data suggested GC Tfh cells were entirely refractory to transcriptional change over the first three days of re-infection, Tcm/Tfh like cells exhibited a modest non-proliferative response, while Th1 memory cells mounted a rapid, dynamic response marked by early IFN-γ production and subsequent proliferation.

### Th1 recall features two waves of RNA processing associated with rapid effector function and subsequent proliferation

Since flow cytometric and scRNA-seq assessment of PbTIIs during re-infection had revealed temporally distinct periods for upregulation of IFN-γ compared to Ki67, we reasoned that a co-ordinated series of transcriptional changes accompanied the progression of Th1-memory cells from a quiescent state prior to re-infection, to effector and proliferative states during re-infection. To map Th1-recall dynamics, we segregated those PbTII recall transcriptomes with prominent Th1 signatures (Figure 3A), and performed pseudo-temporal ordering via Bayesian Gaussian Processes Latent Variable Modelling (BGPLVM) using *GPFates*^13^. This generated a progression of Th1 transcriptomes that correlated with sampled time points (Figure 4A). We then grouped all highly dynamic genes according to their expression dynamics using *SpatialDE*^20^ (Supplementary Table 4). This revealed 9 distinct dynamics, ranging in size from 150 genes – 2314 genes (Figure 4B). Gene Ontology (GO) enrichment analysis indicated some immune-associated genes in Dynamic 1, *e.g. Tcf7, Il7r,* and *Bcl2* that rapidly waned after activation (Figure 4B). These were followed by four prominent waves of transcription, with Dynamics 3 and 5 emerging last, comprised of 886 genes associated with cellular proliferation (Figure 4B and 4C), and preceded by “immune-activation” genes including *Ifng*, *Tnf*, *Il2*, and *Tbx21* in Dynamic 2. Unexpectedly, prior to the peak of “immune-activation” genes was Dynamic 4, its 370 genes strongly enriched in processes related to RNA biology (Figure 4C). These genes were upregulated substantially earlier than cellular proliferation genes during re-infection, but largely overlapped with them during primary infection (Figure 4D). In addition, RNA processing genes overlapped in their dynamics with immune activation genes, raising the hypothesis that genes within these groupings might be transcriptionally linked. Indeed, transcriptional network analysis of Dynamics 2 and 4, specifically during mid-pseudotime, revealed an apparent linking of the two (Figure 4E), with several transcription factors implicated (*Irf8, Irf4, Foxp1, Zeb2, Stat5a, Apex1, Cnbp* and *Fubp1*), as well as *Il2ra* and *Tnfsf8* (encoding CD30L) (Figure 4E). This link was not observed at latter points in pseudotime (Extended Data Fig. 4A), consistent with a transient transcriptional link between RNA processing and immune effector function, which subsequently switched to a link with cellular proliferation (Extended Data Fig. 4B). Together, our dynamic analysis revealed two waves of RNA processing, firstly linked to rapid immune effector function, and later associated with cellular proliferation (Figure 4B). Further examination of genes within Dynamic 4 identified three RNA-associated pathways: ribosome biogenesis^21^ and splicing pathways^22^, which are involved in post-transcriptional/translational processes, and in addition, polyamine metabolism, previously reported to influence CD4^+^ T cell fate during primary immune responses^23^ (Figure 4F). Together our data highlight that in contrast to primary Th1 responses, Th1 recall is characterised by two waves of RNA processing, in particular an early burst that is transcriptionally linked to immune effector function. Furthermore, our data suggest that during recall, genes controlling cellular proliferation are transcriptionally unlinked to those mediating rapid effector function.

**Figure 4:**
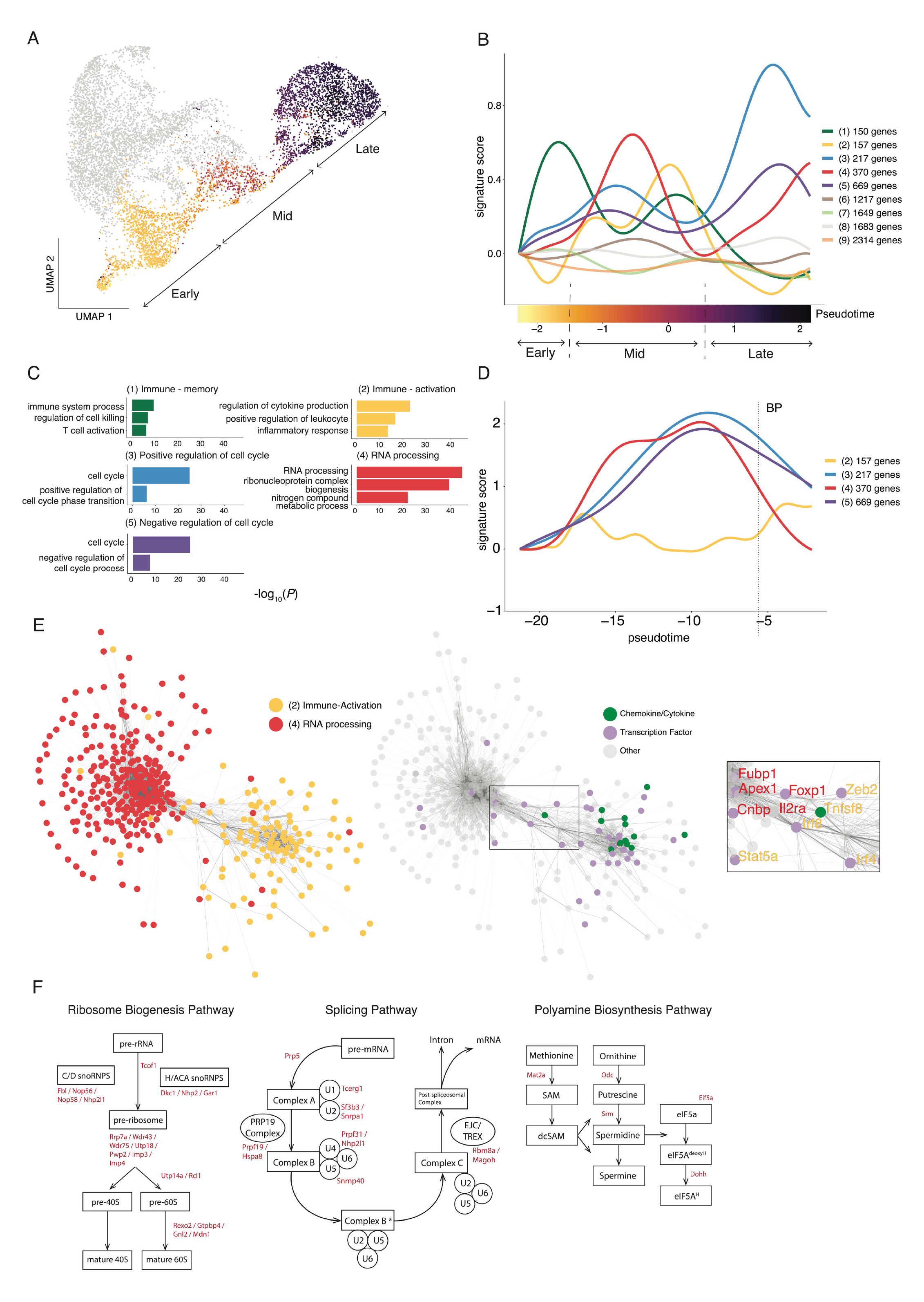
Transcriptome dynamics of Th1 recall reveals two waves of RNA processing associated with rapid effector function and subsequent proliferation. **(A)** UMAP of antigen-experienced PbTII cells prior to and 1, 2, and 3 days post-re-infection, with only Th1-like cells coloured according to inferred pseudotime values (from 1-dimensional bGPLVM, *GPfates*), split into early, mid and late-pseudotime. **(B)** Expression dynamics for pseudo-temporally variable genes in Th1-like cells along pseudotime, genes grouped according to similar dynamics, represented as signature scores. **(C)** Summary of gene ontology enrichment analysis of biological processes associated with genes for selected groups from **(B)**. **(D)** Expression dynamics for gene groups 2, 3, 4, and 5 for for Th1-like PbTII cells during primary infection^14^; BP denotes Th1/Tfh bifurcation point. **(E)** Co-expression network analysis of genes (represented as nodes) in dynamics 2 and 4. Edge weight corresponds to Spearman’s rho values (rho > 0.2). Gene labels in the inset coloured according to dynamic of origin. **(F)** Schematics of enriched biological pathways associated with genes from dynamic 4. Genes found in dynamic 4 highlighted in red.

### Th1 effector recall is characterised by upregulation of a select few genes

We next searched for genes uniquely expressed during Th1-recall compared to primary Th1 responses. We computationally integrated Th1-recall transcriptomes across pseudotime with those from naïve and primary Th1 PbTIIs from our previous study^14^, using *single-cell Variational Inference* (*scVI*) to account for batch effects^24^. UMAP visualisation suggested naïve cells and Th1-memory cells prior to re-infection were transcriptionally distinct from each other, and from primary Th1 cells (Figure 5A). Importantly, Th1-recall transcriptomes at later stages of recall largely overlapped with primary Th1, suggesting substantial similarity between the two, particularly at the peak of Th1 activity. This was corroborated by a relative lack (49 genes) of uniquely upregulated genes in Th1-recall compared to primary Th1 (both compared to naïve PbTIIs) (Figure 5B). GO term enrichment analysis revealed the 49 uniquely upregulated genes were associated with cell cycle (Figure 5B). Instead, Th1-recall upregulated genes (compared to naïve) were essentially a smaller subset (548 out of 1319) of those upregulated in primary Th1 (Figure 5B). Together, our data support the view that no unique genes marked Th1-recall over primary Th1 responses, and moreover that Th1-recall represented a trimmed down version of primary responses.

**Figure 5:**
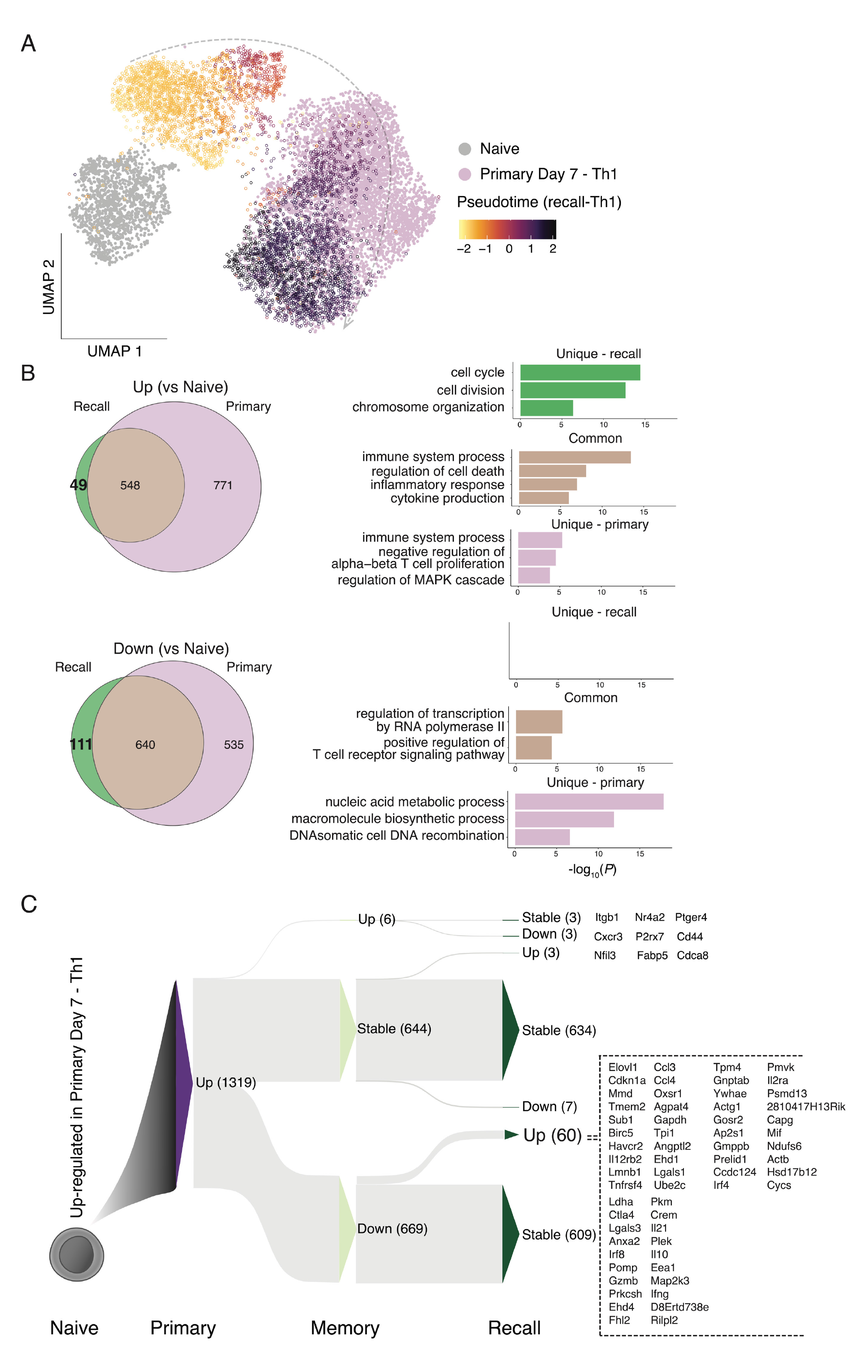
Th1 recall is characterised by upregulation of a select few genes. **(A)** UMAP of PbTII cells after *scVI* integration of naïve and primary Th1 scRNA-seq data from Soon et al.^14^ with Th1-recall scRNA-seq in this study (open circles) coloured according to pseudotime values (after 1-dimensional bGPLVM) – apparent Th1-recall trajectory indicated with arrow. **(B)** (Left) Venn diagrams showing overlap in lists of differentially expressed genes (bayes factor > 3) from peak Th1-primary (relative to naïve PbTIIs), compared to peak Th1-recall (relative to naïve PbTIIs). (Right) Summary of gene ontology enrichment analysis of biological processes associated with genes common or unique to each comparison. **(C)** Sankey plot schematic summarising dynamics of 1319 genes initially upregulated in Th1-primary cells (compared to naïve PbTIIs), as cells proceed to memory and then to peak Th1-recall (Up: upregulated, Down: downregulated, Stable: no significant change; at each stage in reference to the preceding stage). Full list of all 60 genes upregulated from Th1-memory to recall shown on the right.

Finally, we consolidated in a single map the transcriptional path as primary Th1 cells transitioned to Th1 memory and recall (1319 upregulated genes in primary Th1 cells shown in Figure 5C and 1175 downregulated genes in primary Th1 cells shown in Extended Data Fig. 5). Out of 1319 upregulated genes, approximately 50% reverted to naïve levels in Th1 memory, leaving 634 genes stably upregulated in memory and recall (Figure 5C; Supplementary Table 5). While immunological roles for many of these genes, including *Cxcr6* and *Tbx21* are well established at least during primary responses, others such as *Myo1f* and *Maf* remained to be studied in memory CD4^+^ T cells *in vivo*. Crucially, during Th1-recall only 4.8% (63/1319) of genes were further upregulated compared to quiescent Th1-memory cells (Figure 5C). These included secreted molecules, *e.g. Ifng, Il10, Il21, Ccl3, Ccl4* and *Gzmb*, costimulatory markers or receptors (*Havcr2, Ctla4, Il2ra, Il12rb2, Tnfrsf4*) and transcription factors *Irf4* and *Irf8* (Figure 5C). Taken together, these data indicate that Th1-recall is not characterised by transcriptional upregulation of hundreds or thousands of genes as observed during primary infection, but instead of a small gene set highly enriched in chemokine-, cytokine-, and co-stimulation-associated markers.

### TCR diverse, antigen-experienced CD4^+^ T cells in the spleen also exhibit varied propensity for recall during re-infection

Using TCR transgenic PbTIIs, we had noted varying magnitudes and dynamics of transcriptional responses mounted by splenic antigen-experienced CD4^+^ T cells during re-infection. To determine the relevance of our findings to TCR diverse polyclonal CD4^+^ T cells, we first transferred naïve PbTIIs, infected and drug-cured mice as above, and determined if co-expression of the markers CD11a, as previously reported^25, 26, 27^, and CXCR3 (as suggested from our previous study^14, 28^) would permit flow-cytometric enrichment of polyclonal, activated CD4^+^ T cells (Figure 6A). We noted firstly, that PbTIIs prior to and 2 days after re-infection expressed high levels of CD11a and CXCR3 (Figure 6A). Similarly, CD11a^lo^ CXCR3^lo^ polyclonal cells exhibited no upregulation of canonical Tfh/Th1 markers CXCR5 or CXCR6, while comparator CD11a^hi^ CXCR3^hi^ cells expressed high levels of both (Figure 6A). These suggested that flow-cytometric sorting of based on CD11a/CXCR3 would allow for enrichment of *Plasmodium-*specific antigen-experienced CD4^+^ T cells.

**Figure 6:**
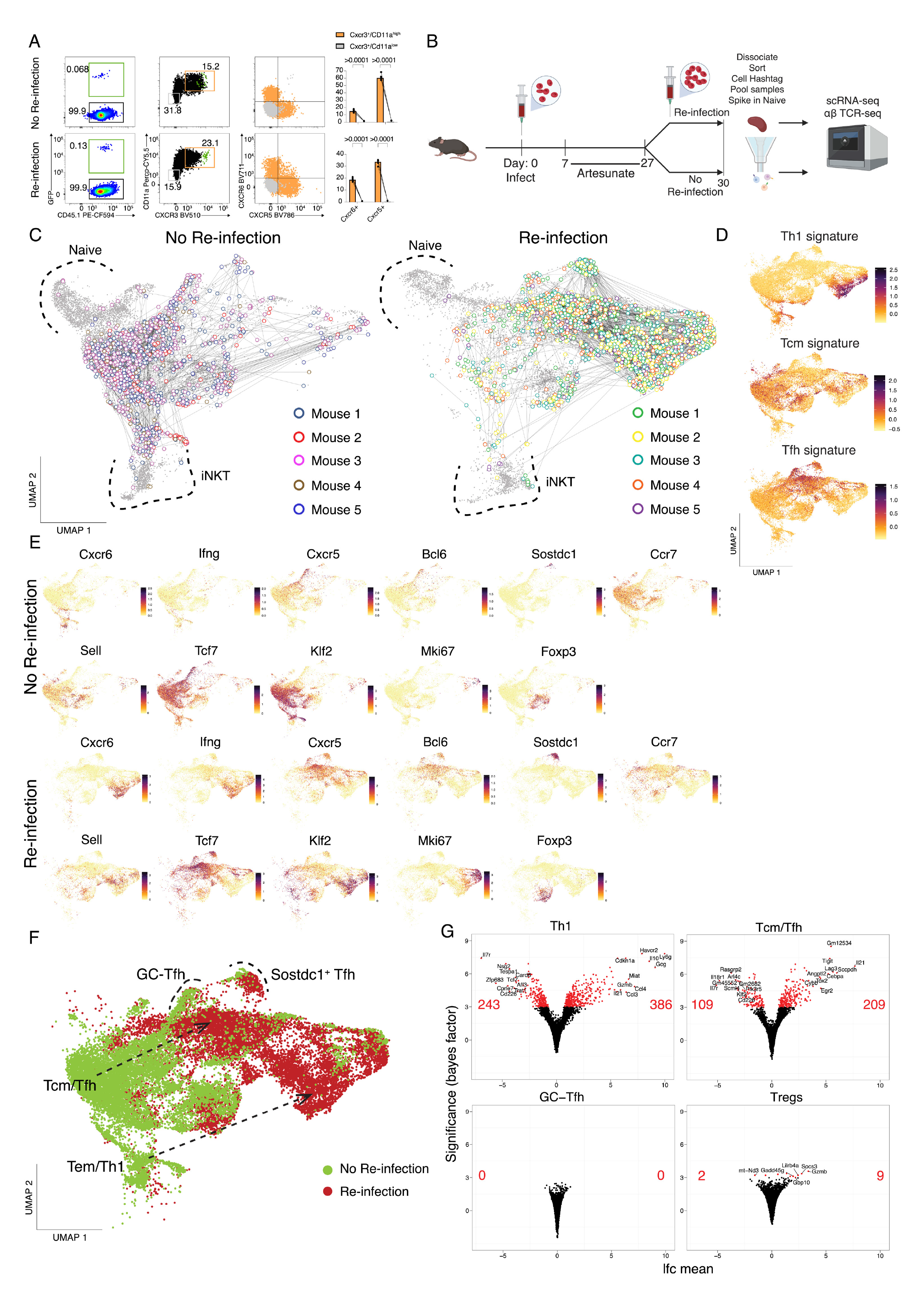
TCR-diverse, antigen-experienced CD4^+^ T cells exhibit varied recall responses during re-infection. **(A)** Representative FACS plots showing GFP^+^ PbTII cells, CD11a and CXCR3 expression, and CXCR6 and CXCR5 expression (orange: gated on CD11a^high^/CXCR3^+^, grey: gated on CD11a^low^/CXCR3^-^) in CD4^+^ T cells prior to and 3 days after re-infection. Graphs show the percentage of CXCR6^+^ and CXCR5^+^ cells from CD11a^high^/CXCR3^+^ versus CD11a^low^/CXCR3^-^ CD4^+^ T cells (n=5 mice). Statistical analysis was performed using a Wilcoxon signed-ranks test. ****p<0.0001. **(B)** Schematic of scRNA-seq + TCR-seq experiment to study CD11a^high^/CXCR3^+^ polyclonal CD4^+^ T cells prior to and 3 days after re-infection. **(C)** UMAP of CD11a^high^/CXCR3^+^ and inferred naïve (CD11a^low^/CXCR3^-^) polyclonal CD4^+^ T cells. Cells sharing the same TCR chains connected with edges. Only families with four or more cells sharing the same TCR chains are shown on the plot. Naïve cells and iNKT cells marked with dotted boundaries. **(D)** UMAP of CD11a^high^/CXCR3^+^ cells expressing Th1, Tcm, and Tfh signature scores. **(E)** UMAP of CD11a^high^/CXCR3^+^ cells expressing genes associated with Th1 (*Cxcr6*, *Ifng*), Tfh (*Cxcr5*, *Bcl6*), Tcm (*Ccr7*, *Sell*, *Tcf7*, *Klf2*), proliferation (*Mki67*), *Sostdc1*^+^ cells, and Tregs (*Foxp3*^+^) cells. **(F)** UMAP of CD11a^high^/CXCR3^+^ polyclonal CD4^+^ T cells prior to and 3 days post re-infection. Differentiation trajectories indicated with arrows. GC Tfh cells and *Sostdc1*^+^ Tfh cells marked with dotted boundaries. **(G)** Volcano plots showing the number of differentially expressed genes (genes with bayes factor > 3) comparing prior to and after re-infection for each inferred state: Th1 cells (Top-Left), Tcm/Tfh cells (Top-Right), GC Tfh cells (Bottom-Left), and *Foxp3*^+^ cells (Bottom-Right). Number of significantly upregulated/downregulated genes and the top 10 upregulated/downregulated genes annotated on volcano plots.

Next, we performed droplet-based scRNA-seq and VDJ sequencing on CD11a^hi^ CXCR3^hi^ polyclonal CD4^+^ T cells recovered from the spleens of individual mice prior to, and 3 days after re-infection, with a 5% spike-in of CD11a^lo^ CXCR3^lo^ control cells (Figure 6B). After de-multiplexing, removal of doublets and low-quality transcriptomes (Extended Data Fig. 6A), TCR genes were removed from the gene expression matrix (Extended Data Fig. 6B) and contaminating non-CD4^+^ T cells were removed (Extended Data Fig. 6C). We also inferred the presence of naïve CD4^+^ T cells based on frequency, lack of clonal connections, and low expression of *Cxcr3* (Extended Data Fig. 6D), as well as noting a minor contaminant of invariant Natural Killer T cells (iNKT), based on expression of semi-invariant TCR chains and *Zbtb16* (Extended Data Fig. 6D)^29^. UMAP visualisation prior to and 3-days after re-infection revealed heterogeneous populations of polyclonal CD4^+^ T cells (Figure 6C) with substantial sharing of TCR chains within and across certain transcriptomically distinct clusters in every mouse (Figure 6C and Extended Data Fig. 7). Having removed inferred naïve and iNKT cells, we noted specific areas enriched for Th1, Tfh and Tcm-like gene expression signatures (Figure 6D) and canonical genes (Figure 6E). We also noted, as for PbTIIs, populations of GC Tfh cells expressing high levels of *Cxcr5* and *Bcl6*, and a neighbouring distinct cluster expressing high levels of *Sostdc1* (Figure 6E). Importantly, prior to re-infection, Tcm and Tfh-like transcriptomes appeared as a continuum, with Th1-like transcriptomes exhibiting a degree of separation, thus replicating our observations with PbTIIs (Figure 6E and F). In contrast to PbTII data, there was a *Foxp3*-expressing cluster with limited clonal connections to other cell-states, consistent with rare instances of iTreg differentiation during primary infection (Figure 6C and 6E).

**Figure 7:**
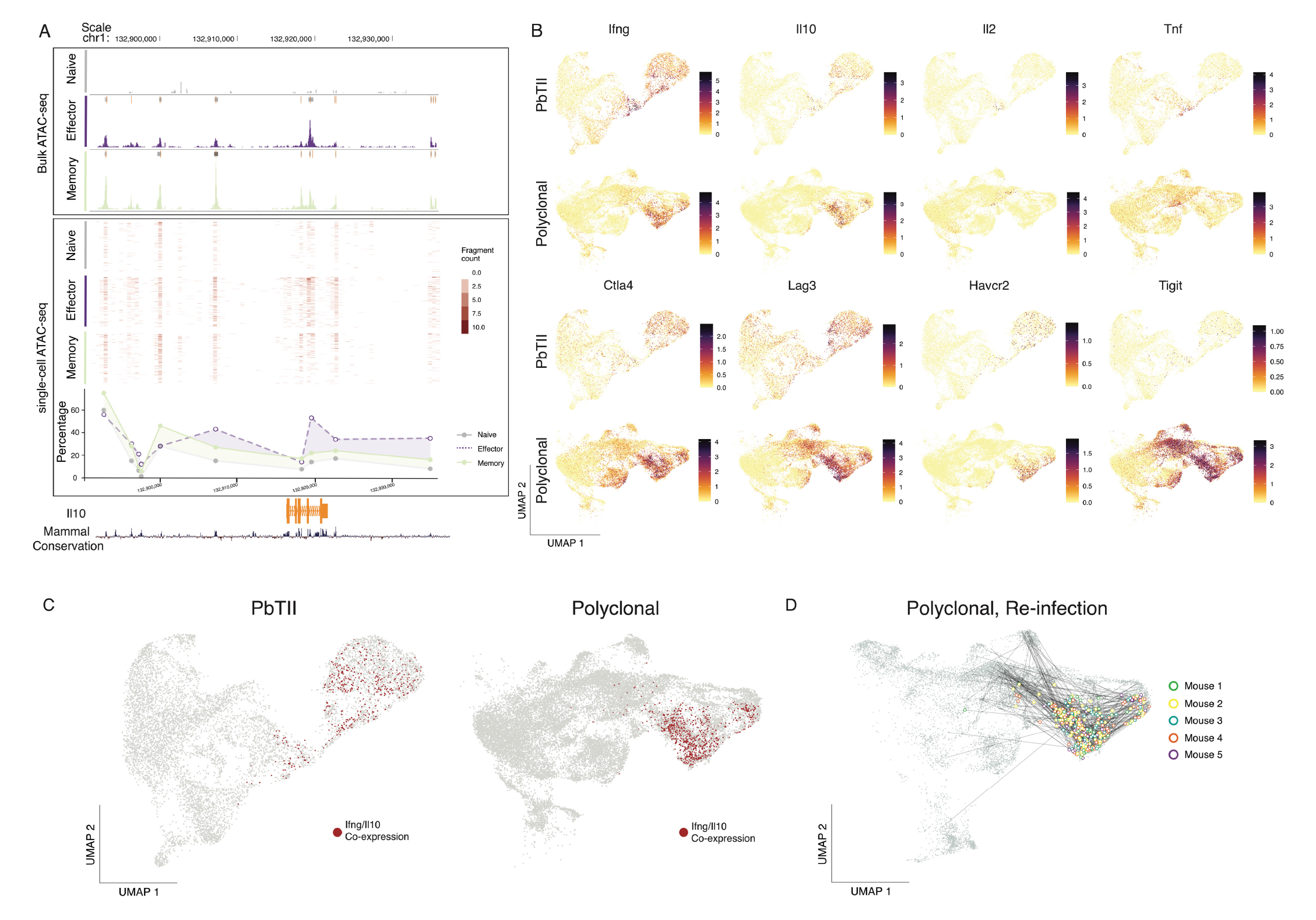
Th1 memory clones exhibit a robust proliferative Tr1 recall response during re-infection. **(A)** UCSC genome browser tracks displaying genomic accessibility signals around the *Il10* gene locus. (Top) Mean bulk ATAC-seq genome coverage. Boxes at the top of each coverage represent peaks called using MACS2. Data representative of two independent experiments showing similar results^14^. (Middle) Accessibility profiles of 200 randomly selected single cells from plate-based single-cell ATAC-seq data^14^. (Bottom) Percentage of cells with open accessibility at each genomic region **(B)** UMAP of PbTII cells and polyclonal CD4^+^ T cells showing mRNA expression of *Ifng*, *Il10*, *Il2*, *Tnf*, *Ctla4*, *Lag3*, *Havcr2*, and *Tigit*. **(C)** UMAP of PbTII cells (Left) and polyclonal CD4^+^ T cells (Right) with *Il10* and *Ifng* co-expressing cells highlighted in red. **(D)** UMAP of CD11a^high^/CXCR3^+^ polyclonal CD4^+^ T cells (with 5% spike-in of CD11a^low^/CXCR3^-^ naïve cells) showing TCR sharing (straight edges) from those cells co-expressing *Il10* and *Ifng* (highlighted in open circles and coloured by mouse of origin; n=5 mice).

3-days after re-infection, CD11a^hi^ CXCR3^hi^ CD4^+^ T cells continued to exhibit heterogeneous states that were clonally related to each other, both within and between clusters (Figure 6C). In particular, we noted strong gene expression signatures for Th1, Tfh, GC Tfh and *Sostdc1^+^* GC Tfh, as well as rarer clonal links with *Foxp3-*expressing cells (Figure 6C and 6D). These data suggest that naïve CD4^+^ T cells gave rise to multiple cell-states during recall, within individual clones populating Th1, Tfh and *Sostdc1^+^* Tfh states. Crucially, comparison of transcriptomic states prior to and after re-infection suggested, as with PbTIIs, that Th1 cells changed substantially while GC Tfh states remained unaltered (Figure 6F), although the relative frequency of *Sostdc1^+^* to *Sostdc1^-^* GC Tfh cells changed after re-infection (Figure 6E and 6F). Based on proximity within UMAPs, we inferred those cells with the strongest Tcm phenotype prior to re-infection, likely adopted a moderate Tfh-like signature after re-infection, with upregulation of *Cxcr5* and *Bcl6* and loss of *Ccr7, Sell, Klf2* (Figure 6E and 6F). Finally, the number of differentially expressed genes for each inferred cell-type prior to versus 3-days after re-infection was substantially higher for Th1-memory cells than Tcm-like cells, with GC Tfh cells being entirely refractory to change over this time-period. Tregs also showed limited transcriptional change during recall (Figure 6G; Supplementary Table 6). Thus, similar to PbTIIs (Figure 6G; Supplementary Tables 7-9), we concluded that GC Tfh cells of diverse specificities remain largely refractory during re-infection with malaria parasites.

### Th1-memory clones exhibit a robust proliferative Tr1 recall response during re-infection

Given that multiple exposures to *Plasmodium* parasites in children have been associated with a switch within antigen-experienced CD4^+^ T cells from the production of pro-inflammatory cytokines TNF, IFN-γ and IL-2 to immunoregulatory IL-10 by a Tr1 state^6^, we sought to specifically examine Tr1 transcriptional dynamics during re-infection in mice. Firstly, analysis of chromatin accessibility via bulk ATAC-seq of PbTIIs, generated in our previous study^14^, suggested increased accessibility around the *Il10* locus at memory stages compared to naïve states (Figure 7A). This was corroborated by plate-based single-cell ATAC-seq, in which a greater proportion of PbTII cells displayed accessibility around the *Il10* locus at memory stages than at naïve (Figure 7A). Together, these data suggested *Il10* might be heterogeneously upregulated by CD4^+^ T cells during re-infection. Indeed, we noted both PbTII cells and polyclonal CD4^+^ T cells displayed robust and prolonged co-expression of *Il10* and *Ifng* during re-infection, consistent with their definition as Tr1 cells in malaria (Figure 7B and 7C). In contrast, *Il2 and Tnf* were only transiently upregulated by PbTII cells during re-infection, (Figure 7B). Furthermore, both PbTII and polyclonal CD4^+^ T cells upregulated immune checkpoint or inhibitory molecules, *Ctla4, Lag3, Havcr2 and Tigit* upon recall (Figure 7B). Thus, our data suggested that primary infection triggers Th1-memory cells that do not sustain TNF or IL-2 production upon re-infection, and instead exhibit a prolonged Tr1 recall response. Furthermore, TCR clonal analysis of polyclonal CD4^+^ T cells suggested clonal links of Tr1 cells to other cell states including Tcm/Tfh-like cells and *Sostdc1^+^* GC Tfh (Figure 7G). Therefore, single naïve CD4^+^ T cells gave rise to multiple states during recall including *Sostdc1^+^* GC Tfh that remain refractory during re-infection, and Tr1 cells that readily re-express IFN-γ and IL-10.

## Discussion

In this study, we sought to understand how antigen-specific CD4^+^ T cells, activated and positioned in the spleen after experimental malaria in mice, responded thereafter to a second infection. In doing so, we aimed to model the experiences of children living in malaria-endemic regions, who are exposed to sequential infections over short periods of time. Although some of their circulating CD4^+^ T cells appear to become increasingly immunoregulatory over multiple exposures (as inferred from antigenic re-stimulation *in vitro* leading to IL-10 production^6^), how this relates to all antigen-experienced CD4^+^ T cells in secondary lymphoid tissues remains unclear. Moreover, given that CD4^+^ T-cell dependent, isotype-switched, affinity-matured IgG is a cardinal mechanism of adaptive immunity to malaria^30^, the effect of subsequent infection on ongoing Tfh and other CD4^+^ T cell responses is unknown.

In our mouse model, a complex spectrum of transcriptional states and micro-anatomical locations was observed in CD4^+^ T cells after primary infection, even for those of a single specificity. This highlights that acquisition of multiple states by a single T-cell clone is common in this model. Whether certain TCR sequences pre-dispose towards GC Tfh or Th1-memory remains to be determined, but based on other model systems^31, 32, 33, 34^, it seems likely that the strength and duration of interactions with professional and non-professional antigen-presenting cells plays a role in giving rise to such a complex landscape of GC Tfh, Th1-memory and other fate choices.

During re-infection, which elicited a transient but readily controlled pathogen load, we noted unexpectedly different recall responses by the heterogeneous, antigen-experienced CD4^+^ T cells, even when T cells were of the same antigen-specificity, located within the same organ of the animal, in either similar or distinct microanatomical regions. For example, Tcm-like and Th1-memory cells were both located in splenic T-cell zones, and yet displayed dramatically different responses, with Tcm-like cells, but not Th1-memory cells failing to proliferate over a three-day period. This suggests a fundamental difference between Th1-memory and Tcm-like cells in their ability to respond to antigenic stimulation *in vivo*. It is remarkable, that Tcm-like cells, considered a prime mediator of immunological memory, did not appear to enter a proliferative phase during the period of assessment, nor mounted an immune effector response. However, it is also possible that our scRNA-seq-based trajectory inference was unable to track rapid change from Tcm-like to a Th1/Tr1 recall. Alternatively, a longer period of analysis may have been necessary to observe proliferation of Tcm-like cells. For polyclonal Tcm-like cells, there appeared to be an emergence of a Tfh-like phenotype during re-infection, that again was devoid of proliferation. Hence, we hypothesise that Tcm-like cells may indeed be memory Tfh cells, poised to re-express *Bcl6* and *Cxcr5*, and facilitate early interactions with B cells during re-infection.

Perhaps most strikingly, GC Tfh cells appeared completely refractory to transcriptional change for three days during re-infection, neither proliferating nor exhibiting an immune effector recall response. The persistence of GC Tfh four weeks after the initiation and subsequent control of infection (with antimalarial drug treatment), suggests their continued role in affinity maturation of antibodies, which are known to accumulate in circulation over this time-period^12, 35, 36, 37^. Hence, we reasoned that these GC Tfh were functional at the time when a second infection was experienced. The lack of transcriptional change in these cells could be due to numerous, non-mutually exclusive factors. Firstly, GC Tfh might already have been maximally stimulated with antigen provided to them by cognate GC B cells. Secondly, GC Tfh might be sequestered away from a bolus of antigen arriving in the spleen after intravenous injection. Thirdly, GC Tfh may be inherently refractory to proliferation. Future experiments, examining GC Tfh *in vitro* may shed light on their lack of response *in vivo.* It should be noted that although others have variously reported on the capacity of GC Tfh cells to proliferate during an immune response, these studies have focussed on primary immune responses only^38, 39^, not on responses during re-infection. Further studies are warranted on the nature of persisting GC Tfh responses during secondary immune responses.

In stark contrast to GC Tfh cells, we observed that rapid and dynamic changes occurred to Th1-memory cells during re-infection. As expected, Th1-memory cells initiated a wave of immune effector molecule production, including IFN-γ and IL-10, prior to a transcriptional program of cellular proliferation. This confirms, in our system that Th1-memory cells exert function prior to dividing. This raises the question of why a Th1-memory cell need proliferate at all, if immune function has already been initiated. One hypothesis is that Th1 memory cells have evolved the dual features of rapid function and amplification of their responses through division. Given that cellular proliferation requires a set period of time, one might speculate on the benefit of immune function being initiated first, to be followed later by proliferation. In addition, our scRNA-seq based, temporally-informed transcriptional network analysis indicated that no transcriptional links existed between the transcription factor hubs that controlled effector function, versus proliferation. Hence, we propose that during re-infection, the processes controlling the magnitude or quality of recall have no bearing on the amplification of this process via cellular proliferation, the implication being that whatever response is triggered, Th1-memory cells will be subject to proliferative amplification.

In addition, an unexpected dynamic related to RNA processing emerged prior to effector function in Th1-memory cells. This was striking because the same genes had only been upregulated during primary Th1 differentiation as cells progressed through cell division. Hence, we reason that a burst of ribosome biogenesis, RNA splicing and polyamine synthesis, unrelated to proliferation, uniquely marks Th1-recall prior to the emergence of effector function. Of note, the same RNA processing genes were also upregulated during proliferation of memory Th1 cells, resulting a biphasic wave of RNA processing. Given the apparent transcriptional link between RNA processing biology and effector function, via genes such as *Irf4, Irf8, Stat5a*, and *Il2*, we hypothesise that ribosome biogenesis, splicing and polyamine synthesis likely trigger early immune-associated transcription factors that promote a rapid wave of effector gene transcription and protein translation.

Our study sought to examine the similarities and differences between the peak Th1 response in primary responses *versus* re-infection. By focussing our attention on “peak” effector responses, we excluded any genes involved in cellular proliferation. In doing so, we noted that only ∼60 genes were upregulated upon Th1-recall. This set included many genes associated with primary Th1 responses including *Irf4, Irf8, Havcr2, Tnfsf4, Il2ra, Ctla4, Ifng, Il10, Ccl3, Ccl4,* and *Gzmb.* In addition, ∼600 genes were stably maintained by Th1 memory cells compared to naïve counterparts. Many of these genes have yet to have functions ascribed during either primary or secondary immune responses. Our study revealed no genes were uniquely upregulated in recall compared to primary Th1 responses, and also confirmed that genes used by many immunologists to measure recall, including OX40, CD25, and the cytokine IFN-γ were likely the best choices for examining this phenomenon. An important ramification of our genome-scale analysis is that Th1-recall is a focussed, partial facsimile of primary Th1 responses. Hence, it is likely that whatever pre-dispositions are set within Th1 cells during an initial malaria infection, these are likely to be difficult to shift during re-infection. This raises the hypothesis that primary immune responses may be the primary determinant for the quality of any subsequent responses, and that immune interventions should be made before or as a child experiences their first infections with malaria parasites.

Our dynamic analysis also revealed that TNF and IL-2 production were only transiently expressed during recall. Instead, the Tr1 cytokines IL-10 and IFN-γ were robustly expressed even as the cells began to proliferate. This suggests that the Th1-recall responses observed in our mouse model of repeated malaria infection, are in fact potent Tr1 responses. We reason here that these potent Tr1 proliferative responses serve not only to control parasite numbers, but also to do so without triggering unwanted immune-pathology. Our data are consistent with those of children living in malaria-endemic regions^6^, suggesting that repeated infections trigger the emergence of Th1-memory cells in the spleen that can mount potent, proliferative Tr1 responses soon after re-infection. Chromatin accessibility assessments confirmed increases in accessibility around the *Il10* locus as a result of a single infection. It remains to be determined whether sequential infections continue to alter the epigenomic profile of CD4^+^ T cells, both at the *Il10* locus and elsewhere.

In summary, our data indicate that parasite-specific CD4^+^ T cells acquire a complex spectrum of antigen-experienced states in the spleen during experimental malaria, and that re-infection triggers a variety of different responses in these cells. Of note, those CD4^+^ T cells that support antibody-mediated immunity appeared least affected by re-infection, while those associated with pro-inflammatory and immune-regulatory responses were rapidly called upon and their responses amplified through cellular proliferation. Our data highlight that single TCR clones can be mobilised to simultaneously exhibit a range of different phenotypes and recall functions while being located within the same organ. This reveals that during re-infection with malaria parasites, the immune system had already diversified and allocated skills to expanded progeny from primary infection, while a breadth of TCR-sequences serves to cater for antigenic diversity. Thus, improved adaptive immunity to malaria may reside not only in embracing antigen diversity, but also by sculpting as early as possible the immunological memories preserved in CD4^+^ T cells.

## Methods

### Mice

C57BL/6J were purchased from Animal Resources Centre (Western Australia) and PbTII and PbTII.nzEGFP mice were bred in-house. All mice were maintained under specific pathogen-free conditions within the Biological Research facility of the Doherty institute for Infection and Immunity (Melbourne, Victoria, Australia), or within the Animal Facility at QIMR Berghofer Medical Research Institute (Brisbane, Queensland, Australia). All animals used were females at 6-12 weeks old. All animal procedures were approved by the University of Melbourne Animal Ethics Committee (1915018) and the QIMR Berghofer Medical Research Institute Animal Ethics Committee (approval no. A1503-601M).

### Adoptive transfer

Naïve spleens from PbTII and PbTII.nzEGFP mice were harvested and mashed through 70-100µm cell strainers and the red blood cells (RBCs) were lysed using RBC Lysing Buffer Hybri-Max (Sigma-Aldrich) or BD Pharm Lyse™ (BD). CD4^+^ T cells were enriched via Magnetic-activated Cell Sorting (MACS) using CD4 (L3T4) microbeads (Miltenyi Biotec). In some instances, PbTII cells were labelled with Violet Proliferation Dye 450 (VPD450) (BD biosciences) prior to the adoptive transfer. For that, enriched PbTII cells were washed twice in Dulbecco’s phosphate-buffered saline (D-PBS) and stained with VPD450 at final concentration of 1µM at 37°C for 15 minutes. After incubation, cells were washed in D-PBS and resuspended in cold RPMI containing penicillin and streptomycin (RPMI/PS). Finally, 10^4^ PbTII.nzEGPF cells were injected per recipient mouse into the lateral tail vein prior to the primary infection; 10^4^ CTV labelled PbTII cells were injected into recipients at 27 days after primary infection.

### Infection

Thawed *Pc*AS infected blood stabilites from our biobank were used to infect C57BL/6J single passage mice. *Pc*AS-infected RBCs were obtained from the passage mice and 10^5^ parasitized RBCs (pRBCs) were injected into each recipient mouse via lateral tail vein injection.

### Parasitemia assessment

Parasitemia assessment was carried out as previously described ^12^. Briefly, blood samples were collected from the tail vein into RPMI containing 5U/ml of heparin. Diluted blood samples were, then, stained with Hoechst 33342 (10 µg/ml; Sigma-Aldrich) and Syto84 (5µM; Life technologies) at room temperature, for 30 minutes in the dark. The staining was quenched with 10 times the initial volume using RPMI, and percentage of double fluorescent RBCs was assessed via flow cytometry.

### Anti-malarial drug treatment

Artesunate (Guilin Pharmaceutical, kindly provided by J. Mohrle) was dissolved in 5% sodium bicarbonate solution at 50mg/ml to form sodium artesunate and diluted in 0.9% saline (Baxter) to the final concentration of 5mg/ml. Mice received intraperitoneal injections of 1mg of sodium artesunate twice daily from day 7 to day 9, once daily from day 10 to day 16 and twice weekly from day 17 to day 24 post infection. Mice were also treated with pyrimethamine in the drinking water (70mg/L, Sigma Aldrich) from day 7 until day 24 after infection.

### Flow cytometry

Spleens were harvested in cold RPMI/PS and homogenised through a 70/100µm cell strainers to create single-cell suspensions, followed by RBC lysis using Lysing Buffer Hybri-Max (Sigma-Aldrich) or BD Pharm Lyse™ (BD). Cells were then stained with Zombie Yellow viability dye (Biolegend) or Live/Dead aqua (Invitrogen), followed by Fc receptor block (BD). Finally, cells were stained with titrated panels of monoclonal antibodies (Supplementary Table 10) diluted in PBS containing 1% of FCS and 2mM EDTA and samples were incubated 20 minutes on ice in the dark. For intracellular staining, Foxp3 / Transcription Factor Staining Buffer Set (eBiosciences) was used to fix and permeabilise cells prior to staining with panels of monoclonal antibodies (Supplementary Table 10) for 30 minutes on ice in the dark. Finally, cells were washed and acquired on Fortessa cytometer (BD). All data was analysed using FlowJo (10.8.0).

### Cell sorting and scRNA-seq

*PbTIIs*: Spleens were harvested and homogenized through 100µm cell strainers to create single-cell suspensions. RBCs were lysed using Lysing Buffer Hybri-Max (Sigma-Aldrich). Samples were then washed twice in PBS containing 0.5% BSA and 2mM EDTA, and CD4^+^ T cells were enriched using MACS CD4 (L3T4) microbeads (Miltenyi Biotec). Next, enriched cells were prepared for cell sorting by Fc receptor blocking and stained with titrated monoclonal antibodies (Supplementary Table 10). Finally, cells were washed and re-suspended in PBS 2% BSA containing propidium iodide at dilution of 1:500. PbTII cells (CD4^+^TCRVα2^+^TCRVβ12^+^) were sorted using a BD FACS Aria III. After sorting, PbTIIs were loaded onto the Chromium controller and cDNA sequencing libraries were prepared using Single-cell 3’ reagent kits (10X Genomics).

*Polyclonal CD4^+^ T cells*: Spleens were harvested and homogenized through 70µm cell strainers to create single-cell suspensions. RBCs were lysed using BD Pharm Lyse™ (BD). Next, samples were washed and incubated with Fc receptor blocking antibodies followed by titrated monoclonal antibodies and TotalSeq^TM^-C mouse hashtags (Supplementary Table 10) for 20 minutes on ice each, in the dark. Finally, cells were washed twice, resuspended in PBS 2% BSA containing propidium iodide at dilution of 1:500. For each timepoint, samples from 5 individual mouse replicates were pooled together and live, “naïve” TCRβ^+^CD4^+^CXCR3^-^ CD11a^-^ and “experienced” TCRβ^+^CD4^+^CXCR3^+^CD11a^high^ T cells were sorted using a BD FACS Aria III. After sorting, “naïve” and “experienced” cells were pooled at 1:9 ratio and loaded onto the Chromium controller and cDNA sequencing libraries were prepared using Single-cell 5’ reagent kits (10x Genomics).

## Processing of data

*PbTII dataset:* Cell Ranger v3.0.2 (*“cellranger count”*) was used to process 10x Genomics gene expression FASTQ files with 10x Genomics mouse genome v2020-A as a reference.

*Polyclonal dataset:* Cell Ranger v6.1.1 (*“cellranger multi”*) was used to process 10x Genomics gene expression, VDJ T cell receptor a/b, and mouse-specific hashtag oligos (HTO) FASTQ files with 10x Genomics mouse genome v2020-A and refdata-cellranger-vdj-GRCm38-alts-ensembl-5.0.0 as references.

## Quality control of scRNA-seq data

*PbTII dataset:* Only genes expressed in 3 or more cells were considered. Cells outside the thresholds of 200-5,000 expressed genes and up to 15% mitochondrial content were removed. High-quality transcriptomes of 2672 naïve PbTII cells and 5423 antigen-experienced PbTII cells were considered for analysis shown in Figure 2 (Supplementary Information Fig. 1A). Naïve PbTII cells were marked by lack of *egfp* expression and antigen-experienced PbTII cells were marked by *egfp* expression (Supplementary Information Fig. 1B). From the combined dataset of PbTII cells prior to, and at 1, 2, and 3 days after re-infection, cluster 9 was further identified as a poor-quality cluster and removed from analysis given its low number of detected UMIs and genes and yet appearing as highly proliferative group of cells based on G2M score (Supplementary Information Fig. 2A), further substantiated by integration using *single-cell variational inference* (*scVI*) from scvi-tools package (v0.17.1)^24^, which placed cluster 9 with other highly proliferative cells (clusters 3 and 5) (Supplementary Information Fig. 2B). Altogether high-quality transcriptomes of 9,245 antigen-experienced PbTII cells (marked by *egfp* expression) were proceeded to downstream analysis (Supplementary Information Fig. 2C).

*Polyclonal dataset:* Only genes expressed in 3 or more cells were considered. TCR genes (TRAV-/TRBV-/TRAJ-/TRBJ-) were also removed from the dataset. Cells outside the thresholds of 300-6,000 expressed genes and up to 10% mitochondrial content were removed. Cells identified as doublets were removed by demultiplexing based on HTO using “HTODemux()” function from *Seurat* package (v4.1.0)^15^. Clusters of cells expressing high levels of *Cd74* and *Cd8a* were identified as non-CD4^+^ T cells were removed. A cluster showing enrichment of *Trav11-Traj18* and *Zbtb16* marking iNKT-cells were removed^29^. Altogether high-quality transcriptomes of 28,296 polyclonal cells were proceeded to downstream analysis.

### Normalisation, Feature selection, and Scaling

Normalisation of data, selection of highly variable features, and scaling of data were performed using “SCTransform()” function from *Seurat* package with default settings.

### Dimensionality reduction

Using the Pearson residuals of all highly variable genes computed from *Seurat*’s “SCTransform()”, principal component analysis (PCA) was performed using “RunPCA()” function from *Seurat* package, and the computed principal components (PCs) as inputs to perform uniform manifold approximation and projection (UMAP) using “RunUMAP()” function from *Seurat* package.

### Unsupervised clustering

Unsupervised clustering was performed using “FindNeighbors()” function followed by “FindClusters()” function from *Seurat* package.

### Gene signature scoring

Signature scores were computed for each cell in the dataset using “AddModuleScore()” function from *Seurat* package. The following groups of genes were used as input to generate various signature scores. Th1: *S100a4*, *Cxcr6*, *Nkg7*, *Gzmb*, *Ccr5*, and *Lgals3*, Tfh: *Cxcr5*, *Tox2*, *Asap1*, *Tnfsf8*, *Slamf6*, and *Tbc1d4*, GC Tfh: *Pdcd1*, *Bcl6*, and *S1pr2*, and Tcm: *Sell* and *Ccr7*.

### Data integration

To perform integration of PbTII datasets highlighted in Figure 1 and Extended Data Fig. 1, we used two different approaches: (1) Using *Seurat* package, individual dataset was first processed using “SCTransform()”. Datasets were then merged together and “FindIntegrationAnchors()” function was used to identify anchors across datasets, and finally “IntegrateData()” function was used to integrate data. The integrated assay was used to run PCA and UMAP to generate UMAP cell embeddings representing the integrated space. (2) Using “RunHarmony()” function from *Harmony* package (v0.1.0)^16^, data was integrated, followed by running PCA and UMAP using the *Seurat* package to generate UMAP cell embeddings representing the integrated space. All parameters were kept a default. Each experiment was identified as separate batch.

To perform integration of PbTII datasets highlighted in Supplementary Information Fig. 2 and Figure 5, we used *scVI*. Prior to integration of Th1-recall cells with Th1-like cells from our previous study^14^ highlighted in Figure 5, D0, D7 p.i., and D28 p.i. PbTII cells from our previous study were analysed (Supplementary Information Fig. 3A) and Th1 signature was visualised (Supplementary Information Fig. 3B) to identify and segregate clusters of cells with the highest Th1 signature (Supplementary Information Fig. 3C). *scVI* was also used to integrate PbTII and Polyclonal datasets highlighted in Supplementary Information Fig. 4. All parameters were kept at default except 2 hidden layers were used for encoder and decoder neural networks, negative binomial model used, and number of epochs set to n=1,000,000/total number of cells to train the model. Latent variables were used as input to run UMAP using *Scanpy* package (v1.9.1)^40^.

### Pseudotime inference and Transcriptome dynamics analysis

Pseudotime was inferred based on latent variable 1 coordinates from Bayesian Gaussian Latent Variable Modelling (BGPLVM) dimensionality reduction using *GPFates*^13^. Automatic expression histology (AEH) from *spatialDE* (v1.1.3)^20^ was applied to perform transcriptomic dynamics analysis using parameters number of patterns (*c*) = 9 and lengthscale =0.4, where *c* specifies the number of gene groups, with each group displaying distinct expression dynamic along pseudotime. Only significantly variable genes along pseudotime were considered (FDR < 0.05).

### Differential gene expression analysis

Differential gene expression analysis was performed using “scvi.differential_expression()” function from *scVI*. All expressed genes were considered in generating the model. The following comparisons were done for comparing transcriptomes before and after re-infection: (1) cluster 12 (Th1-memory cells before re-infection) versus cluster 8 (Th1-recall cells after re-infection) from PbTII dataset, (2) cluster 2 (Tcm/Tfh-like cells before re-infection) versus cluster 10 (Tcm/Tfh-like cells after re-infection) from PbTII dataset, (3) cells before re-infection from cluster 9 versus cells after re-infection from cluster 9 (GC Tfh cells) from PbTII dataset, (4) cluster 9 (Th1-memory cells before re-infection) versus cluster 12 (Th1-recall cells after re-infection) from polyclonal dataset, (5) cluster 0 (Tcm-like cells before re-infection) versus cluster 8 (Tcm-like cells after re-infection) from polyclonal dataset, (6) cells before re-infection from cluster 16 versus cells after re-infection from cluster 16 (GC Tfh cells) from polyclonal dataset (Figure 3B for (1-3) and Figure 6G for (4-6)) Clusters from PbTII and polyclonal datasets are shown in Supplementary Information Fig. 4A. Data integration of PbTII and polyclonal datasets was performed to ensure similar transcriptomes were compared in this analysis. Th1, Tcm, and Tfh signatures were visualised on integrated UMAP of PbTII and polyclonal datasets to compare clusters of similar phenotypes (Supplementary Information Fig. 4B). The following comparisons were done to define Th1 response: (1) clusters 5 and 10 (naïve cells) versus cluster 3 (Th1-primary cells) from droplet-based scRNA-seq generated from previous study^14^, (2) clusters 5 and 10 (naïve cells from previous study^14^) versus cluster 8 (Th1-recall cells from PbTII dataset), and (3) cluster 3 (Th1-primary cells) versus cluster 7 (Th1-memory cells) (Figure 5B and 5C). Clusters of PbTII cells from previous study and visualisation Th1 signature in each cluster to compare clusters of similar phenotypes are shown in Supplementary Information Fig. 3C. Genes with bayes_factor > 3 were considered to be differentially expressed.

### Gene ontology term enrichment analysis

Enrichment analysis was performed using *GOrilla* (*p* < 0.05)^41^. Input gene lists were differentially expressed Enriched GO terms were further curated using *REVIGO* to identify unique GO terms based on semantic similarity^42^.

### Spatial transcriptomics

Spatial profiling of transcriptomes from fresh frozen splenic tissue samples were conducted at the Broad Institute using Slide-seqV2^17^. All downstream analyses on spatial transcriptomic datasets were performed as previously described^18^. Briefly, pre-processing and quality control to obtain high quality datasets were followed by deconvolution to estimate cell-type identities and proportional contributions using *cell2location*^43^. Unbiased clustering was performed on neighbourhood-adjusted library size-normalised data to identify germinal centre regions within splenic sections. When plotting CD4 ^+^T cell subsets in Figure 1E and Extended Data Fig. 2, we condensed the colour scales so that any bead with a proportion of these cell types greater than 50% of maximum in the dataset appeared as the highest value in the colour scale. This transformation was not applied when performing cell-cell colocalisation inference.

### Bulk ATAC-seq analysis

Analyses were performed as previously reported^14^. Briefly, sequenced reads were mapped to mouse genome MGSCv37 (mm9). Peak calling and quality control were performed. The resulting bigWig tracks and narrowPeak files were visualised using UCSC genome browser (https://genome.ucsc.edu/).

### Single-cell ATAC-seq analysis

Analyses were performed as previously reported^14^. Briefly, sequenced reads were mapped to mouse genome MGSCv37 (mm9) and peak calling was performed. Quality control was performed to obtain high quality dataset. Aggregate peak coverage was visualised using UCSC genome browser (https://genome.ucsc.edu/). UMAP embeddings were generated from principal components 2-4 following latent semantic analysis of binarized filtered coverage matrix. A gene locus was considered open in accessibility if 1 or more reads were mapped to the region.

---

**Extended Data Fig. 1:**
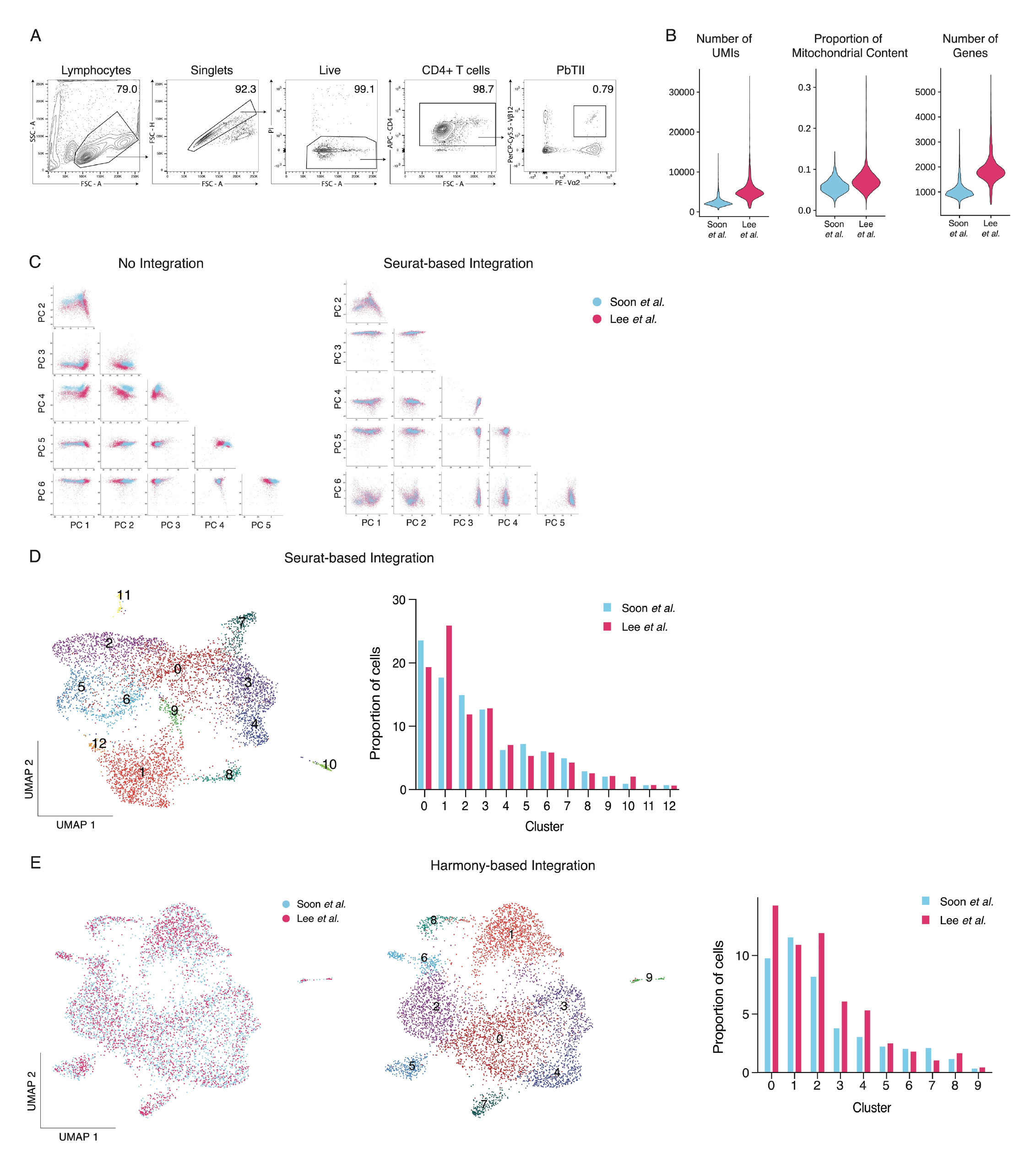
Sorting strategy, quality control and integration of scRNA-seq datasets of PbTII cells 28 days after infection from two independent experiments. **(A)** FACS gating strategy for isolation of PbTII cells from spleen at Day 28 p.i. **(B)** Violin plots showing the distribution of PbTII cells for number of UMIs, proportion of mitochondrial content, and number of genes. **(C)** PCA representation of PbTII cells before and after integration using *Seurat*. Cells coloured by experimental origin. **(D)** (Left) UMAP representation of PbTII cells after integration using *Seurat* showing the individual clusters from unsupervised clustering (Louvain algorithm). (Right) Bar plot showing frequency of PbTII cells from each experiment in each cluster. **(E)** (Left) UMAP representation of PbTII cells after integration using *Harmony*. Cells coloured by experimental origin. (Middle) UMAP representation showing individual clusters from unsupervised clustering (Louvain algorithm). (Right) Bar plot showing the frequency of PbTII cells from each experiment in each cluster.

**Extended Data Fig. 2:**
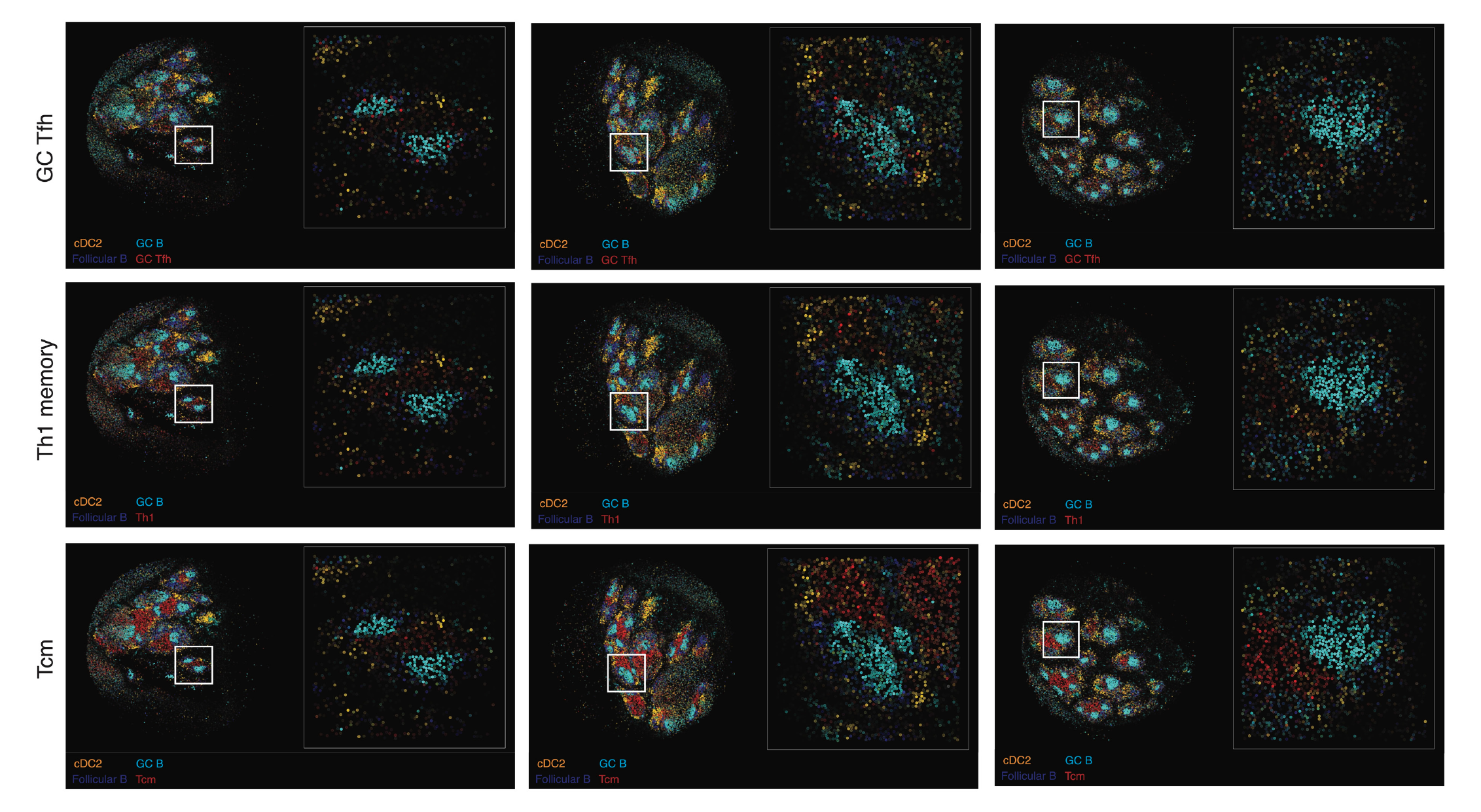
Spatial maps of mouse spleen tissue generated using Slide-seqV2 reveal GC Tfh transcriptomes in GC B-cell regions. Spleens were harvested at day 30 post-infection and processed by *Slide-seqV2* and analysed as in Main Figure 1E. Three independent slides are shown here in three columns, with the same GC B cell, cDC2 and follicular B cell data shown in each panel for a given column, with GC Tfh, Th1 memory or Tcm cells shown in red in each row. These slides are replicates for data shown in Figure 1E.

**Extended Data Fig. 3:**
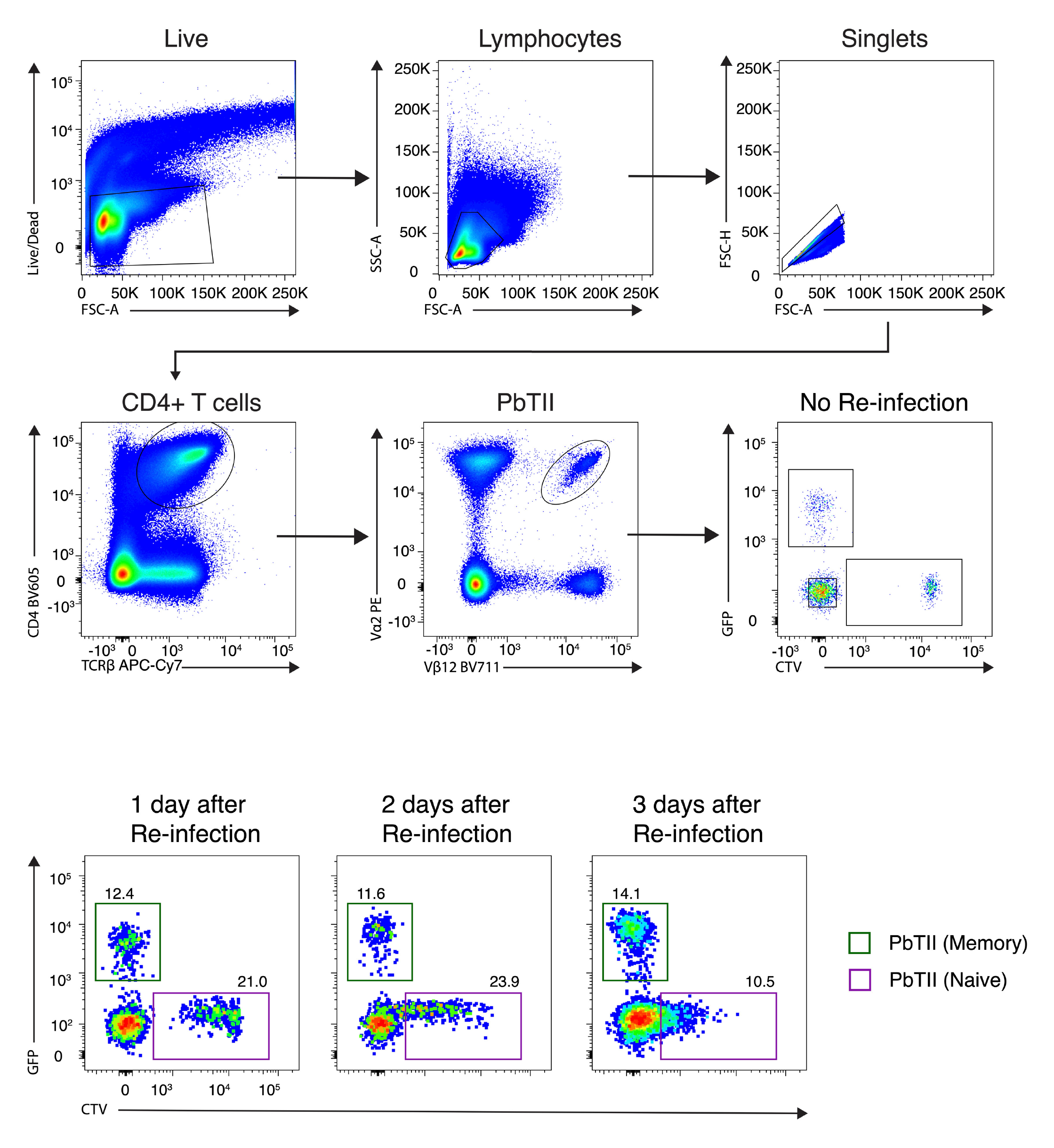
FACS gating strategy for isolation of antigen-experienced and naïve PbTII cells from spleens at prior to (Top) and 1, 2, and 3 days after re-infection (Bottom). These data relate to experiments in Main Figures 1 and 2.

**Extended Data Fig. 4:**
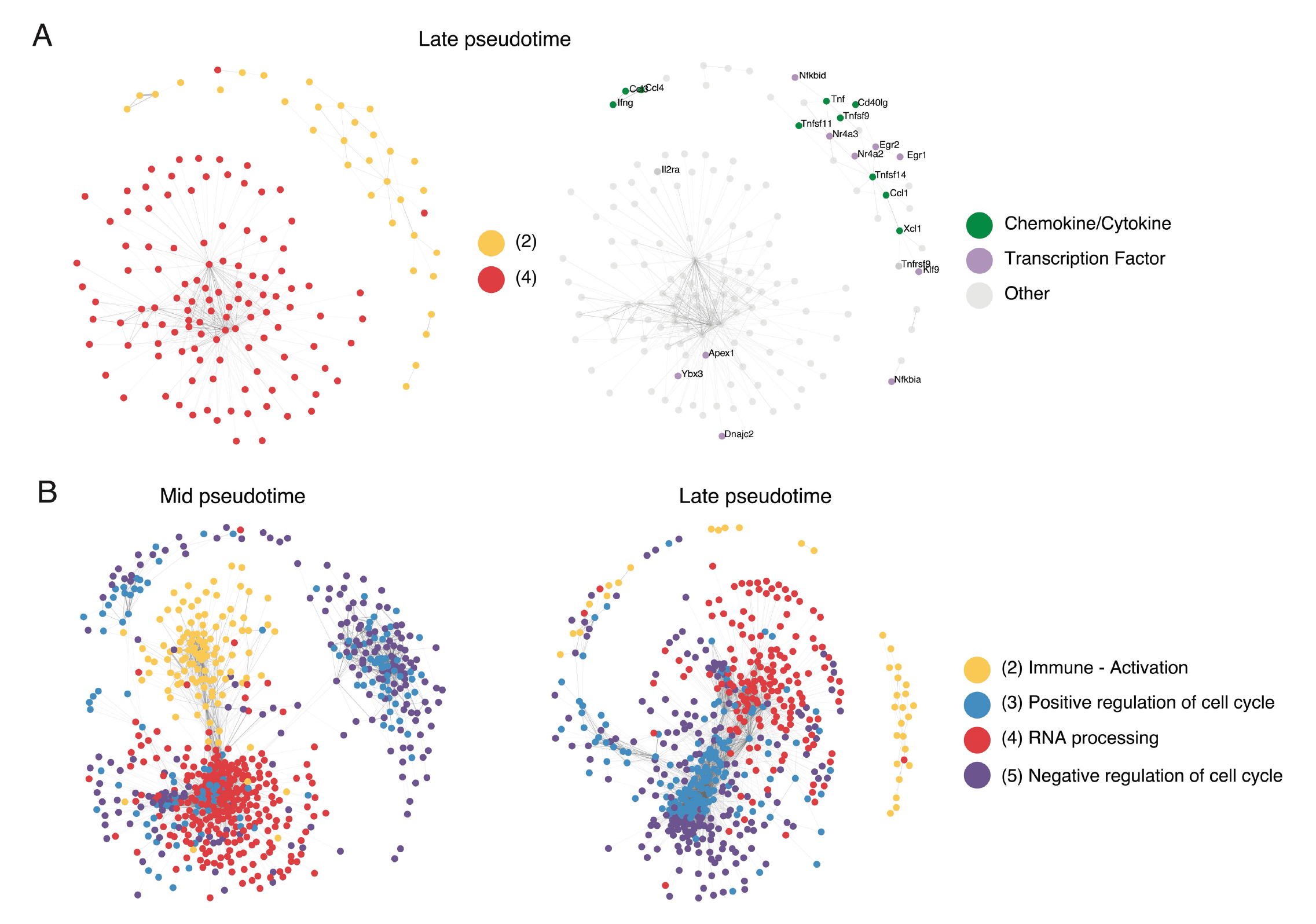
Co-expression gene network analysis shows transient link of RNA processing with immune effector function and later with proliferation. (A) Co-expression network analysis of genes (represented as nodes) in dynamics 2 and 4 using cells from Late pseudotime (in Figure 4A). Nodes are coloured by dynamics of origin (Left) and protein class (Right). (B) Co-expression network analysis of genes in dynamics 2, 3, 4, and 5 using cells from Mid pseudotime (Left) and Late pseudotime (Right). Nodes are coloured by dynamics of origin. Edge weight corresponds to Spearman’s rho values (rho > 0.2).

**Extended Data Fig. 5:**
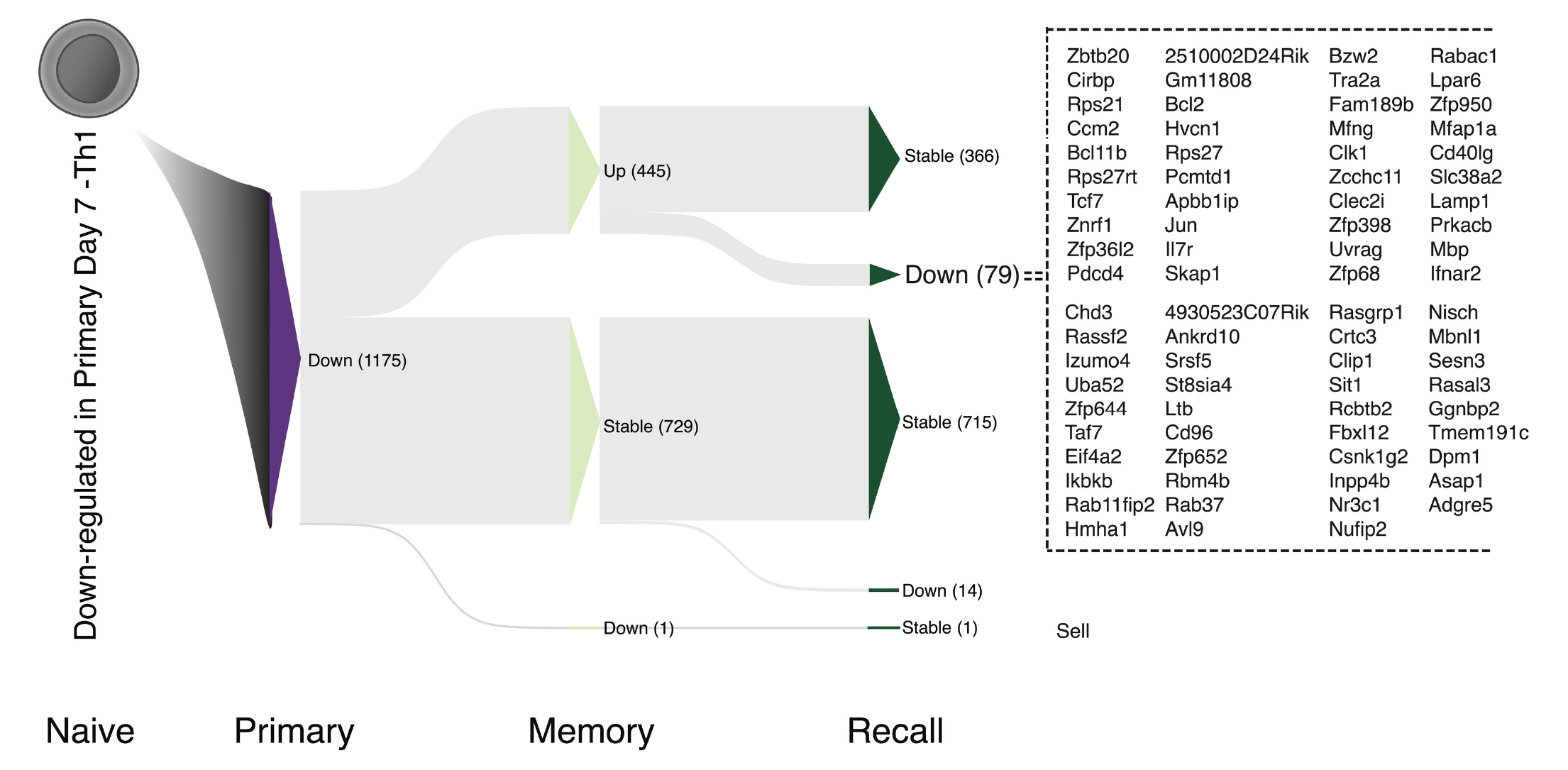
Expression dynamics during memory transition and recall for 1175 genes initially downregulated during primary Th1 differentiation. Sankey plot schematic showing expression dynamics for the 1175 genes initially downregulated in primary Th1 cells relative to naïve PbTIIs (Up: upregulated, Down: downregulated, Stable: no significant change) at each stage in reference to the preceding stage. List of genes upregulated during memory stage and then downregulated during recall shown on the right.

**Extended Data Fig. 6:**
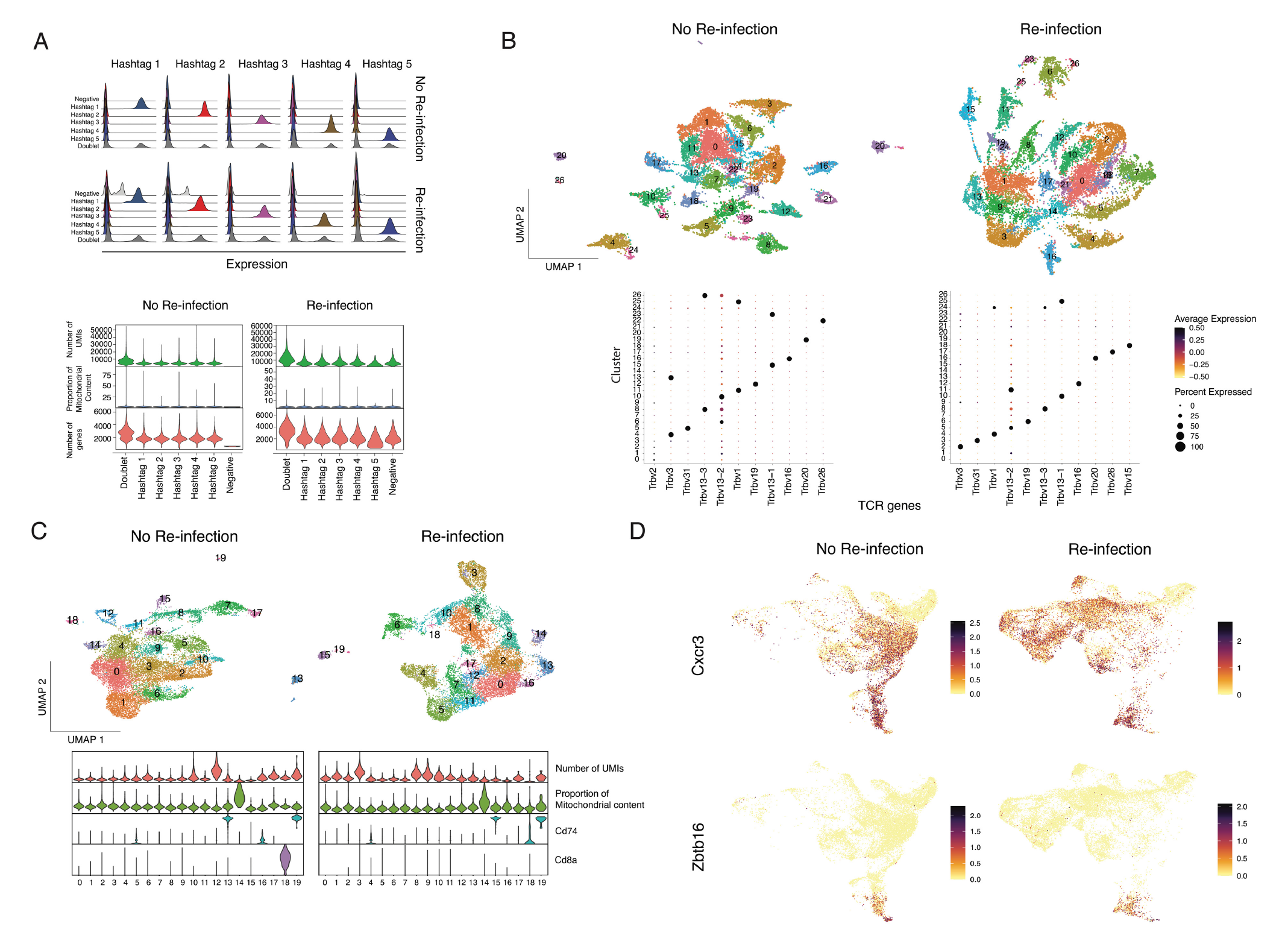
Quality control for scRNA-seq/TCR-seq dataset of CD11a^high^/CXCR3^+^ polyclonal CD4^+^ T cells prior to and 3 days after re-infection. (A) (Top) Ridge plots showing enrichment of hashtag oligos (columns) for negative, singlet (Hashtags 1-5), and doublet groups (rows). (Bottom) Violin plots showing distribution of polyclonal CD4^+^ T cells for number of UMIs, proportion of mitochondrial content, and number of genes detected. (B) (Top) UMAP of polyclonal CD4^+^ T cells prior to and after re-infection before removal of TCR genes. Cells are coloured by clusters from unsupervised clustering (Louvain algorithm). (Bottom) Dot plots showing expression of TCR genes from top 5 marker genes of each cluster. Colour represents mean normalised expression and size of dots represents percentage of cells expressing these genes. (C) (Top) UMAP of polyclonal CD4^+^ T cells after removal of TCR genes. Cells coloured by clusters from unsupervised clustering (Louvain algorithm). (Bottom) Violin plots showing distribution of polyclonal CD4^+^ T cells for number of UMIs, proportion of mitochondrial content, *Cd74* expression, and *Cd8a* expression. (D) UMAP of polyclonal CD4^+^ T cells expressing *Cxcr3* and *Zbtb16* after removal of clusters with high *Cd74* and *Cd8a* expression (shown in (C)).

**Extended Data Fig. 7:**
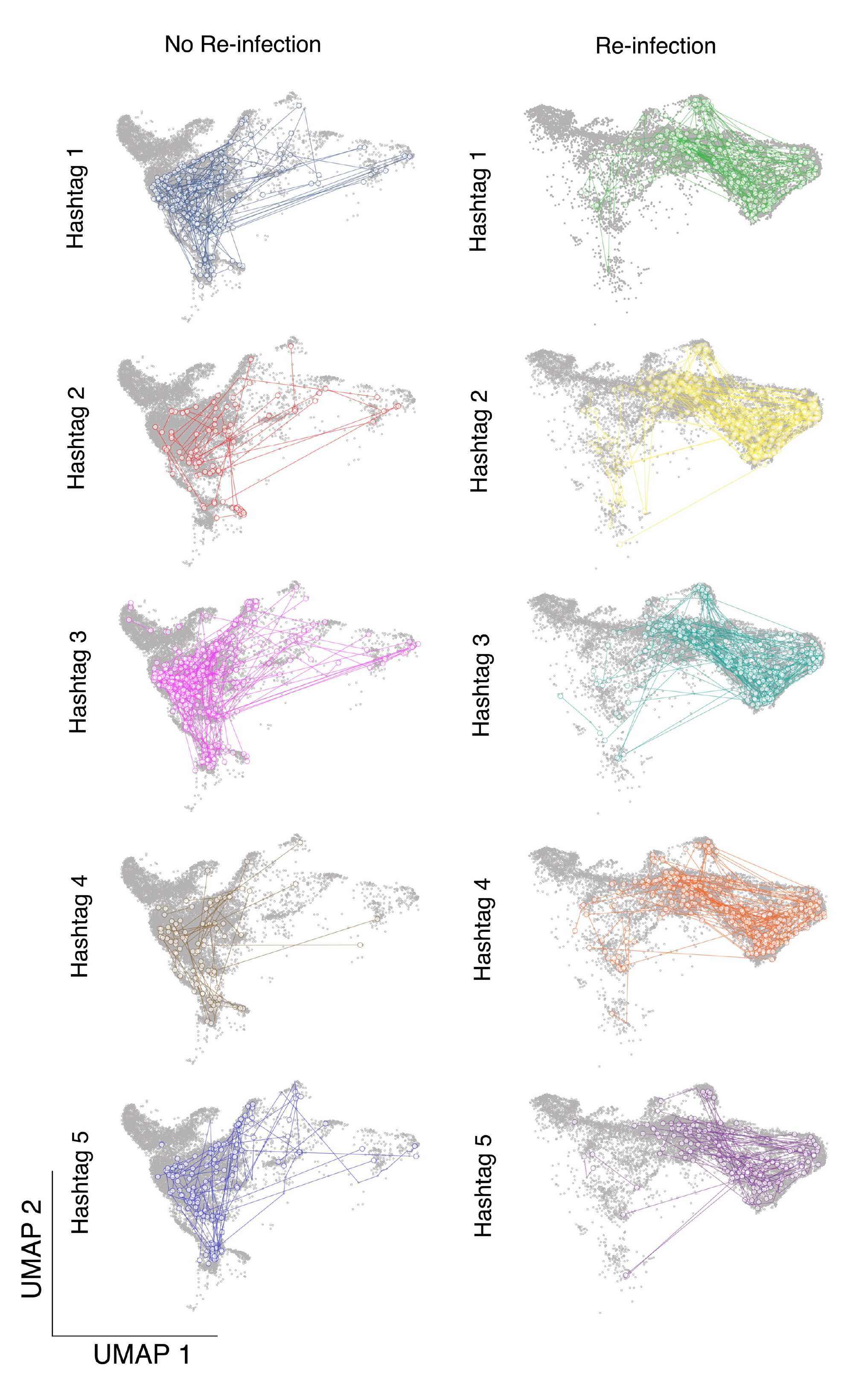
Clonal relationships of polyclonal CD4^+^ T cells prior to and after re-infection. UMAP representation of CD11a^high^/CXCR3^+^ polyclonal CD4^+^ T cells (with 5% spike in of CD11a^low^/CXCR3^-^ naïve cells) for 5 mice prior to re-infection, and 5 mice after re-infection. Cells sharing the same TCR chains are connected by edges and only families with four or more cells sharing the same TCR chains are shown on the plot. Each hashtag represents data from one mouse.

## Notes

### Competing Interest Statement

The authors have declared no competing interest.

