## Supplementary Information for "CD4^+^ T cells display a spectrum of recall dynamics during re-infection with malaria parasites"

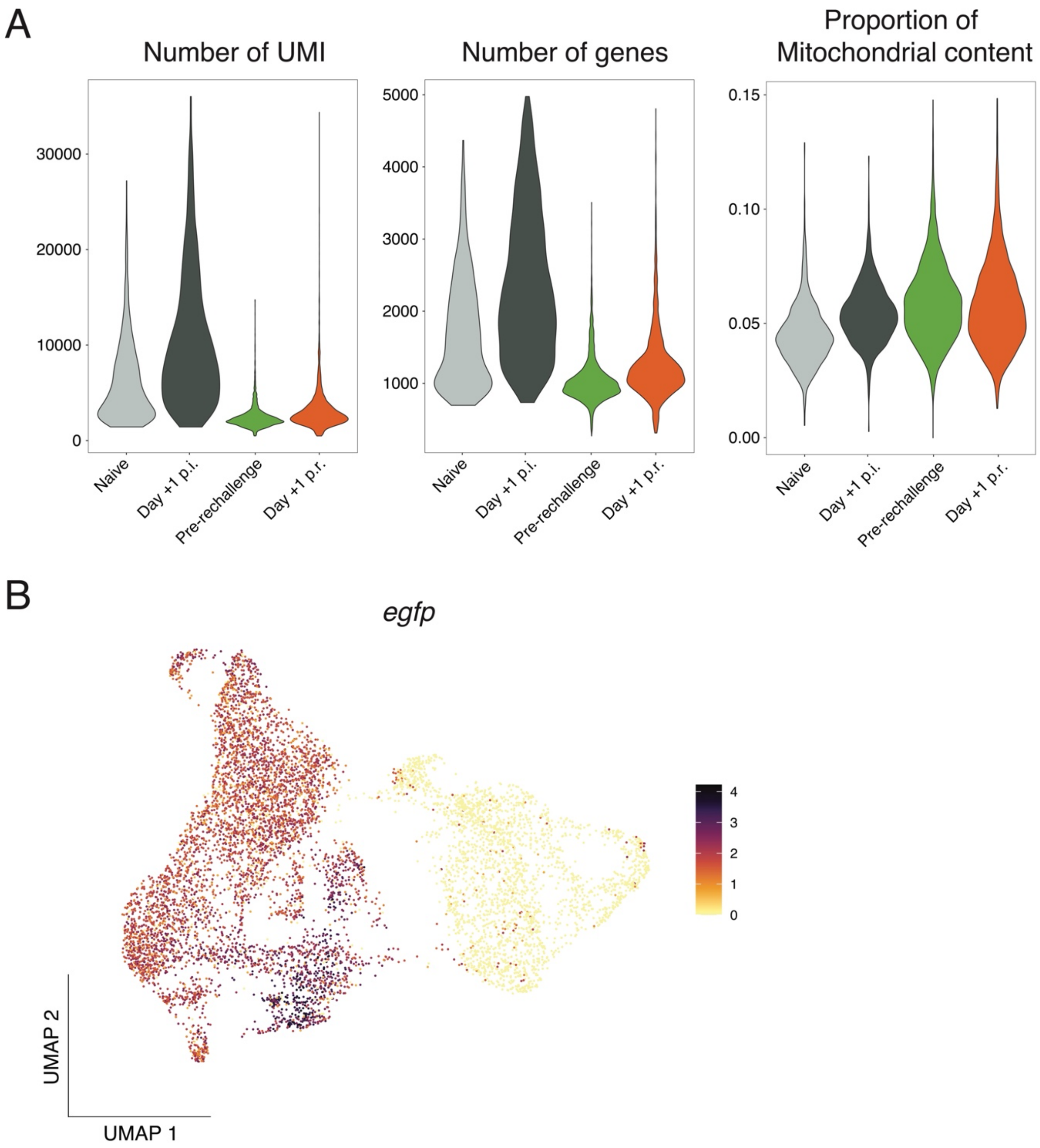


Supplementary Information Fig. 1: **Quality control of scRNA-seq data for antigen-experienced and naïve PbTII cells prior to and 1 day after re-infection (A)** Violin plots showing the distribution of PbTII cells for number of UMIs, proportion of mitochondrial content, and number of genes. **(B)** UMAP visualisation of *egfp* expression, denoting antigen-experienced PbTIIs.

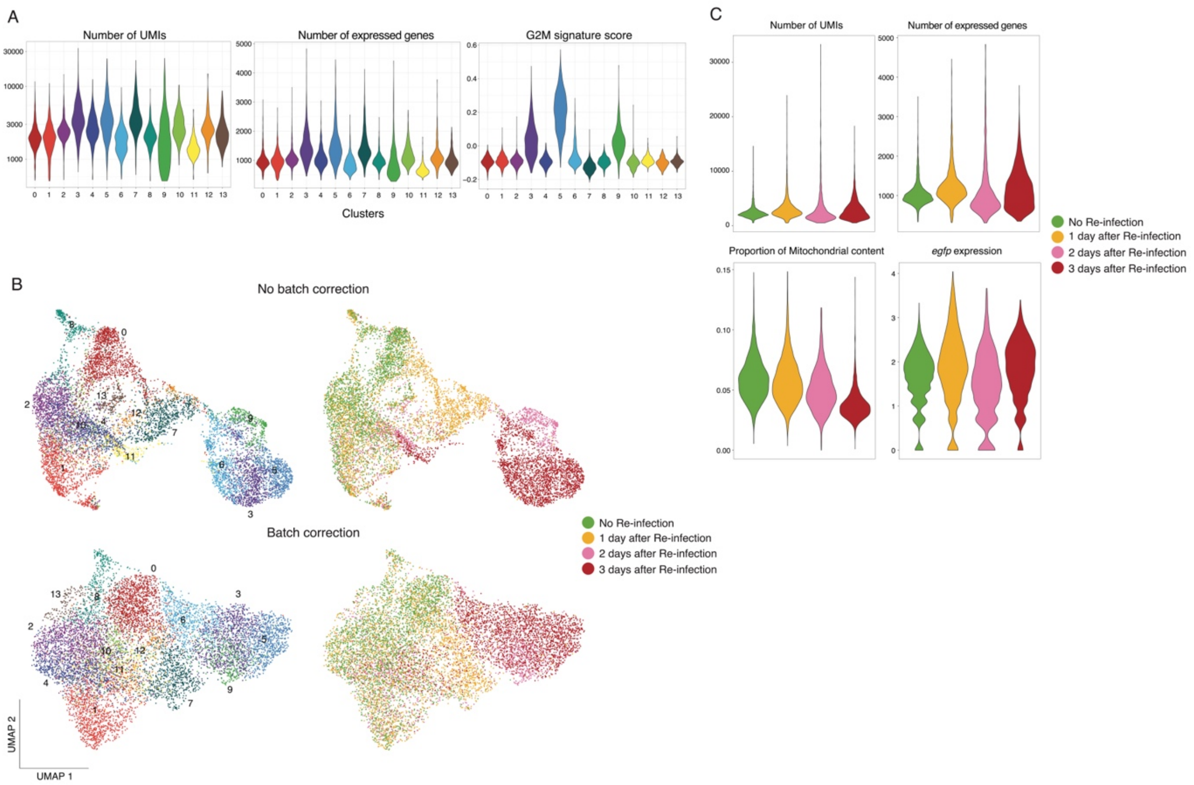


Supplementary Information Fig. 2: **Quality control of scRNA-seq data for PbTII cells prior to and 1, 2, and 3 days after re-infection. (A)** Violin plots showing the distribution of PbTII cells for number of UMIs, number of genes, and G2M signature score for each cluster. **(B)** UMAP representation of PbTII cells before (Top) and after (Bottom) integration using scVI. Cells are coloured by clusters from unsupervised clustering (Leiden algorithm) (Left) and timepoint (Right). **(C)** Violin plots showing the distribution of PbTII cells for number of UMIs, proportion of mitochondrial content, number of genes, *and egfp* expression for each sample after removal of cluster 9 in **(A/B)**.

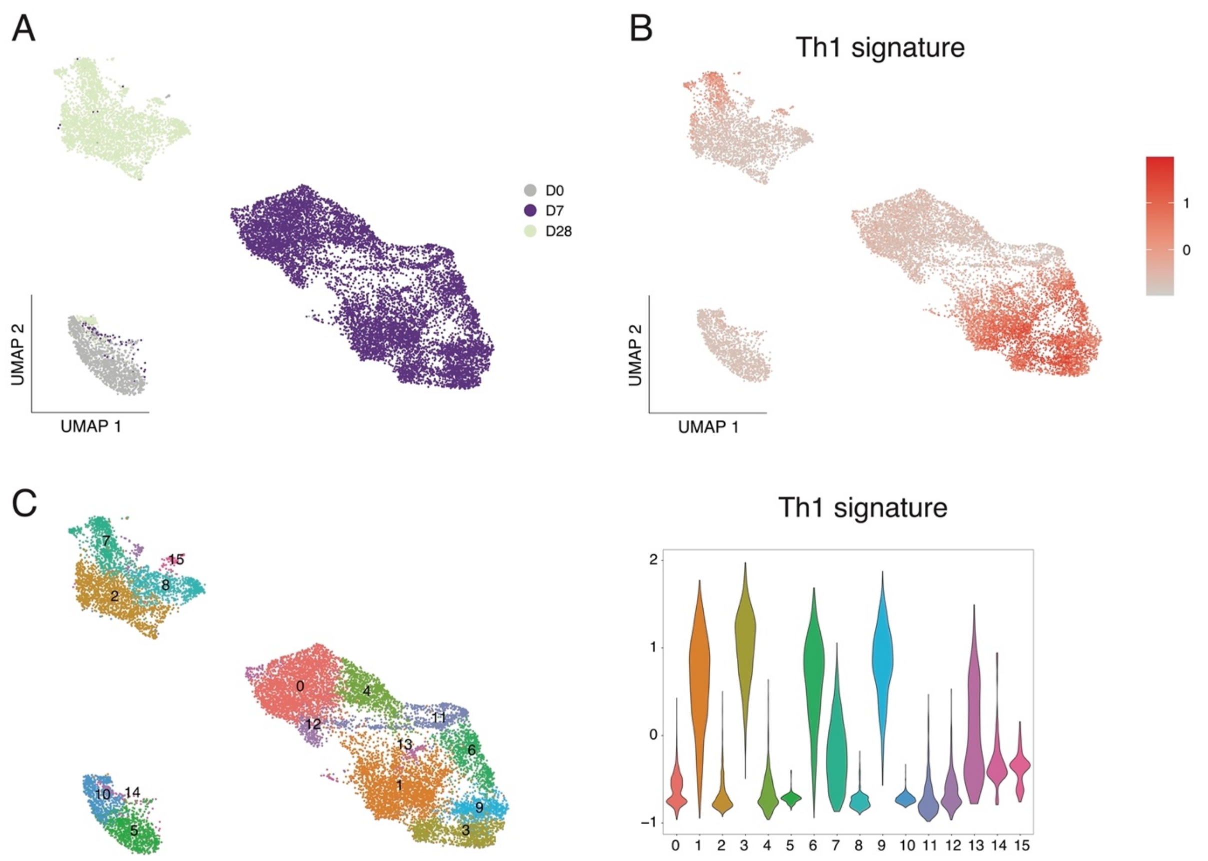


Supplementary Information Fig. 3: **Analysis of scRNA-seq data of naïve and effector PbTII cells at day 7 post-infection, compared with day 28 p.i. (Soon *et al.* 2020). (A)** UMAP representation of PbTII cells. Cells are coloured by sample of origin. **(B)** UMAP visualisation of PbTII cells expressing Th1 signature score. **(C)** (Left) UMAP representation of PbTII cells. Cells are coloured by clusters from unsupervised clustering (Louvain algorithm) and (Right) violin plot showing the distribution of PbTII cells for Th1 signature score for each cluster in **(B)**.

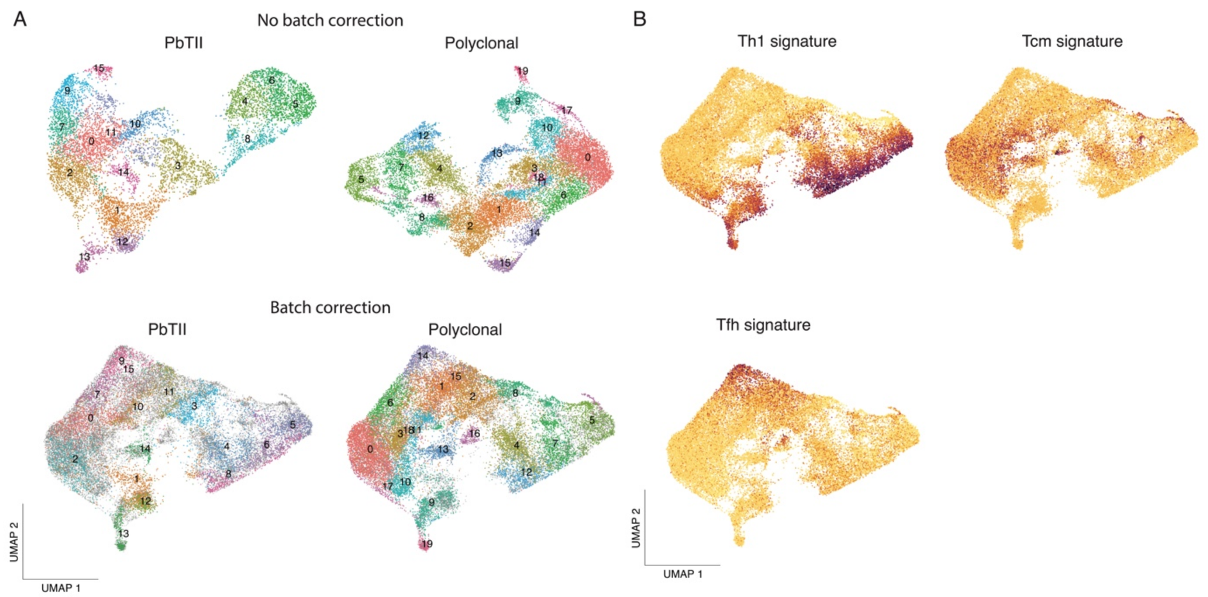


Supplementary Information Fig. 4: **Integration of scRNA-seq datasets of PbTII cells and polyclonal CD4$^+$ T cells prior to and 3 days after re-infection to identify cell clusters of similar phenotypes. (A)** (Top) UMAP representation of PbTII cells and polyclonal CD4$^+$ T cells prior to data integration. Cells are coloured by clusters from unsupervised clustering. (Bottom) Combined and integrated UMAP representation generated using scVI. Cells are coloured by clusters from unsupervised clustering (Louvain algorithm) of PbTII cells (Left) and polyclonal CD4$^+$ T cells (Right). **(B)** Visualisation of Th1, Tcm, and Tfh signature scores on the integrated UMAP shown in (**A** – Bottom).
